# Endolysin B as a new archetype in *M. tuberculosis* treatment

**DOI:** 10.1101/2023.12.05.569960

**Authors:** Anna Griego, Beatrice Antinori, Andrea Spitaleri, Ilaria Muzzolini, Stefano Muzzioli, Luca Calò, Giorgia Moschetti, Marianna Genta, Edoardo Scarpa, Valeria De Matteis, Riccardo Miggiano, Stefano Ricagno, Loris Rizzello

**Author notes:** Corresponding author *Prof. Loris Rizzello, Infection Dynamics Laboratory*. - *Department of Pharmaceutical Science – University of Milan, Via G. Balzaretti 9 – 20133 Milan (IT) National Institute of Molecular Genetics (INGM), Via Francesco Sforza 35 – 20122 Milan (IT) e-mails:.

## Abstract

So far it is thanks to antibiotics that illnesses, such as Tuberculosis (TB), are treatable. However, antimicrobial misuse combined with *Mycobacterium tuberculosis* phenotypic plasticity are nullifying the effects of the existing therapies. As a result, increasingly people are dying (one person dies of TB every 20 seconds), especially due to the rise of multi-and extensively drug-resistant strains. There is indeed a urgent need for new and more effective therapies, which should match the requirement of avoiding the rise in drug resistance. Among them, the use of bacteriophage – bacteria-restricted viruses – or simply phage lytic enzyme (*i.e.,* endolysin) represents one of the most promising alternatives. All phages encode for the endolysin A (LysA), which degrades the bacteria cell wall, finally leading to the release of the newly synthesized virions. Nevertheless, mycobacteriophages (bacterial viruses selectively infecting mycobacteria), evolved the additional endolysin B (LysB) to selectively damage the complex mycobacteria cell wall, and to evade from their host. LysB, owing a lipolytic enzyme, can degrade the thick mycolic acid layer, and hence disrupt the integrity of the mycobacterial membrane. Despite its key role in mediating mycobacteria lysis, the molecular mechanism regulating LysB binding to its target remains poorly characterized. Herein, we selected Ms6LysB and created a fluorescent engineered version as a proxy to analyze LysB binding qualitatively and quantitatively to both the fast-growing non-pathogenic *Mycobacterium smegmatis* and the slow-growing pathogenic *M. tuberculosis.* Additionally, we shed light on LysB antimicrobial activity upon *M. tuberculosis* infection, by using alveolar-like mouse macrophages (mAMs) as a cellular model that closely recapitulates the natural niche of *M. tuberculosis* infection. Our study provides the proof-of-principle that Ms6 LysB binding to the outer mycobacterial membrane can impair *M. tuberculosis* growth homeostasis and that LysB retain its lytic properties even when internalized by mAMs. This lays the groundwork for the use of LysB as a new therapeutic strategy to undermine *M. tuberculosis* infection.

## Introduction

Despite *Mycobacterium tuberculosis’s* ancient origins, this recalcitrant intracellular pathogen continues to extract a heavy toll on global public health. The World Health Organization (WHO) estimated that *M. tuberculosis* still infected 10.3 million individuals and claimed 1.30 million deaths in 2022, remaining, after COVID-19, the world’s second leading cause of death from a single infection agent^1,2^. The exponential appearance of multi-(MDR) and extensively-(XDR) drug-resistant *M. tuberculosis*, only partially controlled by the recent clinical use of Bedaquiline, is further complicating Tuberculosis (TB) treatment and eradication^3,4^. The current chemotherapeutic treatment for *M. tuberculosis* infection requires a prolonged and convoluted antibiotics regimen, which still includes many of the original drugs developed in the 1950s and 1960, whose side effects, together with the length of the treatment, entail poor patient compliance^2,5^. Aside from the rise of MDR and XDR-resistant *M. tuberculosis* mutants due to genetic mutation^6,7^, *M. tuberculosis’* unique thick and waxy cell envelope, together with its slow replication rate, additionally contribute to the bacilli intrinsic phenotypic resistance to most antimicrobials^7,8^. This points out the urgency of conceiving new antitubercular strategies top enhance *M. tuberculosis* clearance, while reducing or avoiding classical antibiotics selective pressure, and hence, the emergence of new drug-resistant strains.

Within the last decade, the administration of mycobacteriophages – viruses that selectively infect and lysate mycobacterial species - has been explored as a new alternative approach to counteract *M. tuberculosis* dissemination^9,10^. However, the low reproducibility in commercial phage preparations, the absence of scientific evidence on phage titer, host range, phage possible immunogenic and side effects, and the absence of proper regulations for its human use resulted in a reduced interest in such approach, especially in the Western countries^11^.

Mycobacteriophages encode within their genome lytic enzymes, namely endolysins, which crucially participate in the release of newly produced viruses by disrupting mycobacterial membrane^9,11^. Because of their intrinsic lytic properties, endolysin has been widely studied as a potential new therapeutic alternative to antibiotics and whole lytic phages for the treatment of bacterial infections^11^. Diversely from other bacteriophages, mycobacteriophages synthesize a second endolysin, endolysin B (LysB), which lipolytic activity disrupts the integrity of the mycolic acid layer^12,13^. Biochemically, LysB hydrolyzes mycolic acid from the mycolylarabinogalactan-peptidoglycan complex and trehalose-6-6’-dimycolate (TDM) from the outer membrane of the mycobacterial cell wall^12,13^. Several experimental observations demonstrated the efficacy of different mycobacteriophage LysBs in diminishing *M. tuberculosis* growth and the low likelihood of developing mycobacterial resistance^14–19^. Yet, the biochemical mechanism behind LysB binding to mycobacterial membrane, and the possibility to retain LysB lytic activity within alveolar macrophages (AMs, the *M. tuberculosis* natural niche^20^), is still unclear.

In this study, binding and activities of Ms6 LysB are thoroughly dissected. We computationally modeled LysB protein and its complex with polysaccharides containing N-acetylglucosamine (GlcNAc), and essential component of the bacterial cell wall. Then, we set up biochemical and flow cytometry assays to quantify LysB affinity for *M. smegmatis* and *M. tuberculosis* membrane, proving LysB’s higher affinity for the pathogenic mycobacteria. Moreover, taking advantage of an *ex vivo* mouse cellular model closely reproducing an alveolar-like phenotype (mAMs)^21,22^, we showed that LysB can abundantly accumulate within mAMs cytoplasm while inducing a pro-inflammatory polarization and the accumulation of lipid droplets within mAMs intracellular environment. Lastly, we analyzed LysB putative antitubercular effect in mAMs infected with *M. tuberculosis*. We demonstrated that although LysB only partially reduced *M. tuberculosis* infection, it can potentially control the infection when AMs are insulted with *M. tuberculosis* as a lower intracellular bacterial burden has been observed.

Hereby, our findings offer a more comprehensive biochemical characterization of LysB binding to *M. tuberculosis* membrane and provide initial evidence of the potential therapeutic benefits of a LysB-based antitubercular therapy.

## Results

### LysB_FL_ diversely binds *M. tuberculosis* and *M. smegmatis* mycomembrane

To quantitatively assess LysB binding to the mycomembrane, we synthesized an inducible bacterial vector expressing LysB C-fused to a superfolder-GFP (sfGFP), here referred to as LysB_FL_. To guarantee LysB and sfGFP independent folding and mobility, a GSGSGS link was added between the two proteins. Parallelly, we synthesized a second inducible vector only carrying the sfGFP, that we used as a control condition. (**Figure S1A**). At first, we optimized and validated both LysB_FL_ and sfGFP expression (**Figures S1B** and **S1C**), and then exploited the presence of a 6X-hist tag at the C-terminal to purify the recombinant proteins (**Figure S1D**). LysB_FL_ protein identity was confirmed by mass spectrometry analysis (**Table 1**). Nex, we used an *M. smegmatis* strains constitutively expressing tdTomato and incubated it with 15 nmol of either LysB_FL_ or sfGFP for 3 hours. After abundantly washing out the unbound proteins, we showed by confocal imaging that, despite previous observation^16,23^, LysB_FL_ can still bind to *M. smegmatis* even in presence of minimal concentration of surfactant adjuvants (0.05% of Twen-80) (**Figure S1D**). Parallelly, by exploiting LysB_FL_ and *M. smegmatis* fluorescence, we quantified the nmol of protein bound to a single bacterium. Data confirmed that the presence of sfGFP do not alter LysB_FL_ intrinsic ability to binds to its target (**Figure S1F**). To further detail the biophysical interactions between LysB_FL_ and the outer membrane (OM) of **(A)** *M. smegmatis*, we used microscale thermophoresis (MST) analyses. Decreasing concentrations of a wild-type strain of *M. smegmatis* were incubated with 1.25 nmol of either purified sfGFP or LysB_FL_ for 3 hours. Although the variation of temperature induced some, albeit minimal, oscillations to the fluorescence of the sfGFP incubated with mycobacteria, we observed a substantial alteration of relative fluorescence when *M. smegmatis* was exposed to LysB_FL_ (**Figure S1G**). To additionally prove LysB_FL_ interaction with *M. smegmatis*, we carried out a Surface Plasmon Resonance (SPR) experiment. Here, two diversely concentrated *M. smegmatis* suspensions (*i.e.* #1, #2) were flowed onto a sensor chip previously functionalized with LysB_FL_ (76.1 RU binding) (**Figure S1H**). Following *M. smegmatis* injection within the sensor chip, we could observe a relevant signal drop, possibly due to buffer mismatch with the mobile phase. Nevertheless, we could also detect an increase in the binding response, qualitatively supporting an interaction event between LysB_FL_ and *M. smegmatis* (**Figure S1H**). Lastly, analysis of the fraction of *M. smegmatis* bacilli positive for LysB_FL_ fluorescence by flow cytometry demonstrated an average of only 39.85 ± 21.5% GFP positive bacteria after 3 hours incubation (**Figure S1I, S1J and S1K**).

Although the mycobacteriophage Ms6 was initially isolate from *M. smegmatis*, it has been shown that Ms6 LysB can also bind to other mycobacterial species, such as the slow-growing pathogenic *M. tuberculosis*^16^. Nonetheless, little is known how LysB_FL_ interacts with *M. tuberculosis* OM. Hence, we incubated 15 nmol of the purified protein, and its relative control (sfGFP), with a strain of *M. tuberculosis* that constitutively expresses tdTomato, and qualitatively assess the LysB_FL_ binding to the *M. tuberculosis* (OM) by high-resolution confocal microscopy (**Figure 1A**). After 3 hours of incubation, we observed the exclusive binding of LysB_FL_ to the OM, suggesting that the sfGFP was not contributing to the interaction^24^. Diversely from previous observations^24^, we demonstrated that the binding of LysB_FL_ to the OM was occurring even in the presence of low concentration of surfactants (0.05% polysorbate 80), which have been generally used to favor the permeation of bacterial membrane and the exposure of the binding motifs^12,16,17,24^. Further quantitative analyses of biding, made by aerosol-free flow cytometry, showed an average of 78.8 ± 30.3% (with a high of 94.3%) of the *M. tuberculosis* population positive to sfGFP after 3 hours of incubation. We imposed an experimental condition where the concentration of LysB_FL_ was constant (15 nmol), irrespective of the number of bacteria initially exposed to the recombinant protein (**Figure 1B, 1C and S1L**).

We verified the accuracy of this data by conducting an *in vitro* binding assay, which relied on the fluorescence intensity ration of the bacterial reporter over the mutant protein. Our results indicated that the concentration of bound protein *per* single bacteria increases as the number of *M. tuberculosis* bacilli increases. We also observed a sharp increment in binding above a threshold of OD_600_ nm equal to 4, possibly indicating that the bacterial concentration might actively influence LysB_FL_ binding efficiency (Figure 1E **and Figure S1M**). Overall, this set of experiments demonstrated that LysB_FL_ can recognize and bind to the OM of *M. tuberculosis*, and the interaction is unaffected by the presence of the fluorescent probe. Finally, we checked whether LysB_FL_ influence *M. tuberculosis* growth. We incubated a planktonic culture of the *M. tuberculosis* strain expressing tdTomato with 15 nmol of either LysB_FL_ or sfGFP and used bacterial fluorescence as a proxy of mycobacterial vitality. We showed that while the control protein only slightly affected *M. tuberculosis* growth over time, LysB_FL_ induced a drop in viability comparable to a high concentration of rifampicin (10x MIC), a first-line antitubercular drug (Figure 1F). This data provides initial indication that LysB_FL_ is not only able to recognize and bind to *M. tuberculosis* OM, but also able to significantly impair the growth of the pathogen. Lastly, by comparing LysB_FL_ the binding towards *M. smegmatis* and *M. tuberculosis*, we showed that LysB_FL_ binding increased, almost linearly, with increasing concentration of bacteria in both mycobacterial species (Figure 1D and **1E**). However, at the highest OD_600nm_ tested, there was a reduction of binding of 0.43 fold compared to *M. tuberculosis*. This suggests that LysB_FL_ owns an intrinsic higher specificity for the outer membrane of *M. tuberculosis*.

**Figure 1.**
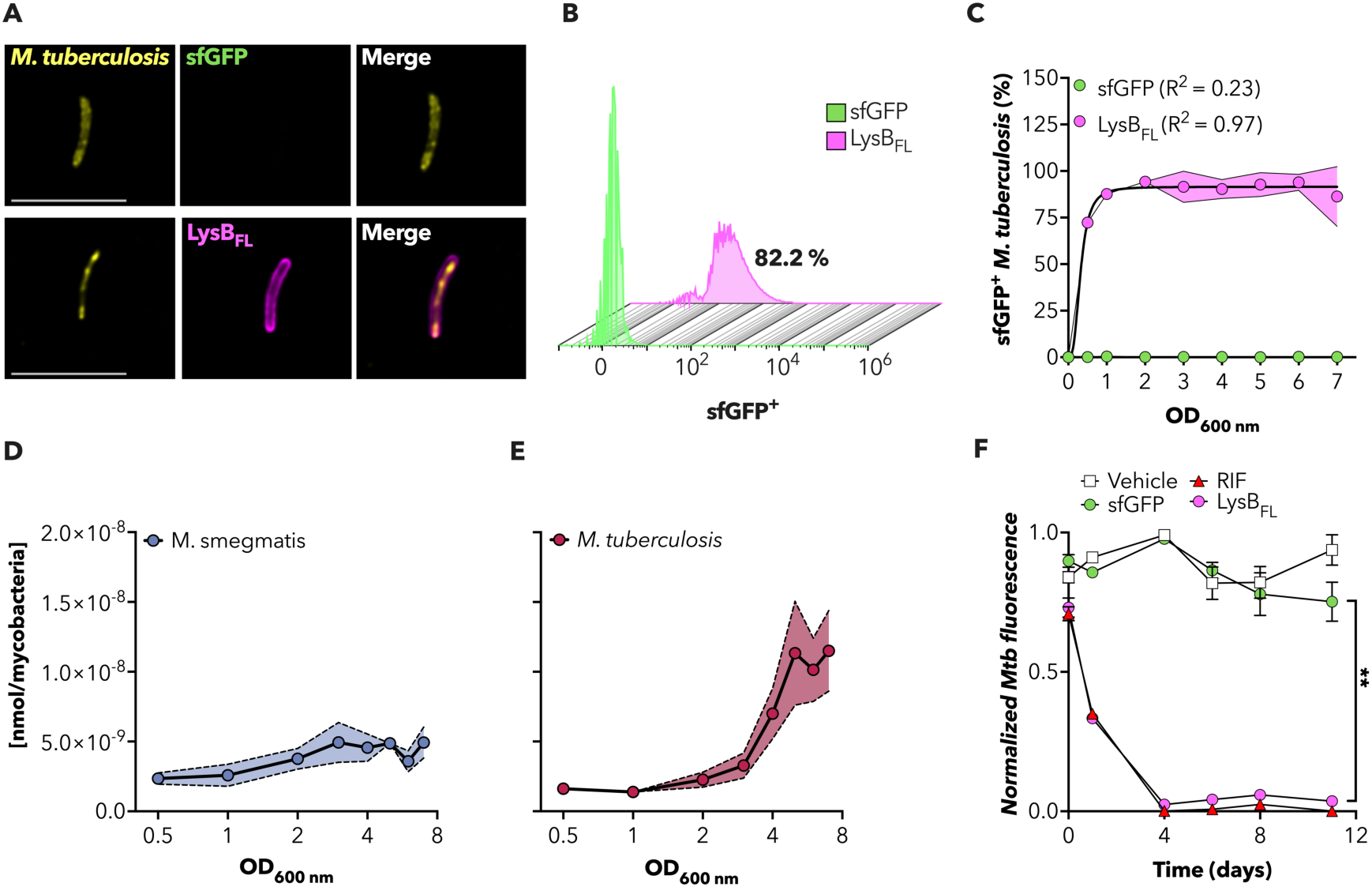
LysB_FL_ diversely binds M. tuberculosis and M. smegmatis mycomembrane. **(A)** Representative images of M. tuberculosis treated for 3 hours with 15 nmol of either sfGFP (upper panel) or LysB_FL_ (lower panel). M. tuberculosis (yellow), sfGFP (green), LysB_FL_ (magenta), are merged. Scale bar 5 μm. **(B)**Histogram showing flow cytometry sfGFP^+^profile detected in the sample with OD_600_ = 6 of bacteria treated with 15 nmol of either sfGFP (green) or LysB_FL_ (magenta) for 3 hours. **(C)**Scatter plot reporting the percentage of sfGFP positive M. tuberculosis bacteria detected by flow cytometry in samples treated for 3 hours with 15 nmol of either sfGFP (green) or LysB_FL_ (magenta). Black lines indicate a Sigmoidal 4PL fitting curve. Magenta shading area denote SD value. The data reported are from two independent experiment. **(D** and **E)** Plate reader assay quantification total nmol of LysB_FL_ bound to M. smegmatis (**D**) or M. tuberculosis (**E**) after 3 hours of incubation with decreasing concentration of mycobacteria. Blue and dark red shading area indicated SD range. The data shown are from 4 independent experiments. **(F)** M. tuberculosis growth kinetics performed measuring tdTomato fluorescence in M. tuberculosis treated with a) the maximum volume of PBS (Vehicle, white square), 10x MIC Rifampicin (Rif, red triangle), 15 nmol of sfGFP (green circle), and 15 nmol of LysB_FL_ (magenta circle). Mean value ± SD (n=3). Asterisks denote significant difference by ONE-way ANOVA: **p = 0.008.

In conclusion, our data confirmed that LysB_FL_ has the intrinsic property of recognizing and binding both *M. smegmatis* and *M. tuberculosis* OM even in the absence of detergent, showing a higher affinity for the OM of the pathogenic species.

### *In silico* molecular characterization of LyB_FL_ binding to the mycomembrane

After quantifying LysB_FL_ binding to mycobacteria OM, we decided to better decipher the molecular force driving such process. Hence, we carried out *in silico* analyses to molecularly determine the interaction between LysB_FL_ and the mycobacterial OM. We used, as a putative LysB_FL_ substrate, 5 repeated units of N-acetylglucosamine (GlcNAc), one of the most abundant polysaccharides present within the mycomembrane^30^, to mimic the OM. We first modeled LysB_FL_ structure by using AlphaFold2 (Figure 2E). We refined our Alphafold2 analysis using long molecular dynamics, finally generating a computational model providing valuable insights into the dynamic behavior and structural stability of the protein (Figure 2A-C). Trajectory data analysis obtained by *in vitro* simulation showed that LysB_FL_ presents a strong persistence of secondary structure and solvent-accessible surface area with respect to the sfGFP protein. The evidence suggest that the inserted linker (SGSGSG) between the sfGFP and LysB_FL_ do not alter either LysB or sfGFP, while guaranteeing the fully accessibility of LysB_FL_ to its target, an important requirement for correct membrane binding assessments. Notably, the *in silico* simulation confirmed the presence of a strong internal correlation within the first about 86 residues of the LysB (Figure 2D). This suggests a high degree of concerted motion and coordination among the atoms in this region. Therefore, the strong internal correlation of the N-terminal region indicates a localized stability, where residues within this segment exhibit synchronized movements. Such coordinated dynamics may point towards a specific functional role or structural motif, which may be crucial for the protein’s stability or interaction with the membrane wall. For instance, it may suggest the presence of critical binding sites, active regions, or domains important for the LysB’s biological activity.

**Figure 2.**
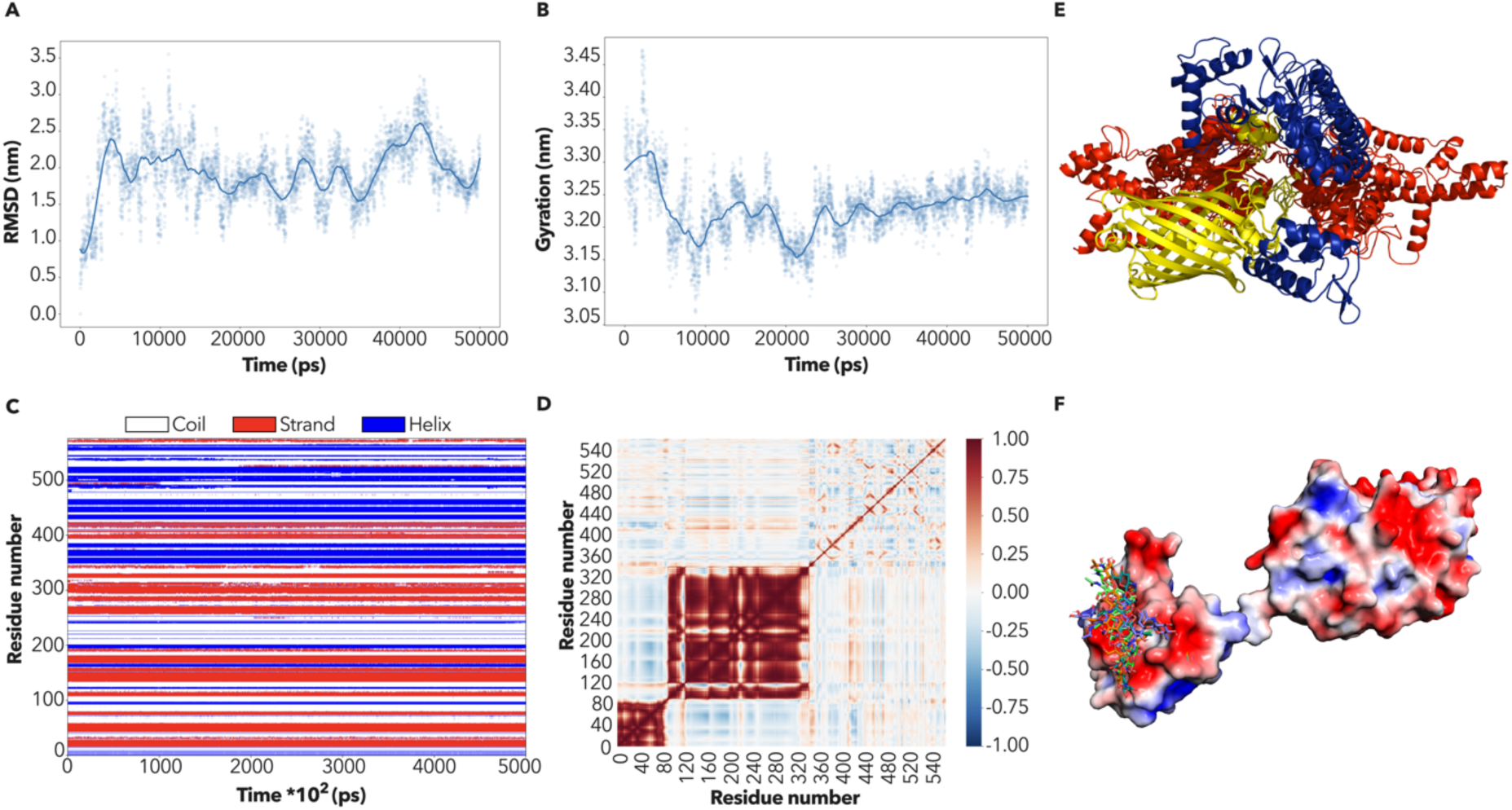
In silico molecular characterization of LyB_FL_ binding to the mycomembrane. **(A)** RMSD time series. **(B)** Gyration radius time series. **(C)** Secondary structure time series. **(D)** Cross correlation matrix from molecular dynamics. **(E)** AlphaFold2 first 4 ranked models overlapped on GFP protein. **(F)** Docking model from HADDOCK 2.4. LysB is shown as surface and colored by electrostatic potential surface. (GlcNAc)_5_ is shown as colored licorices, overlapping the best top 6 cluster coordinates.

We predict the interaction between LysB_FL_ and (GlcNAc)_5_ using HADDOCK2.4 software to provide insights at the atomic level into the binding mode, and the key residues involved in the interaction. The results from the docking calculations indicate the presence of a well-defined and specific binding region on the LysB_FL_ protein for (GlcNAc)_5_. This LysB putative binding site is characterized by a strong negative surface patch, which could make favorable interactions with (GlcNAc)_5_, such as hydrogen bonding, van der Waals forces, and electrostatic interactions, contributing to the stability of the LysB-(GlcNAc)_5_ complex (Figure 2F). The docking results identify specific amino acid residues within the binding site that play a fundamental role in the interaction with (GlcNAc)_5_, including D14, E44, E52, Q54 and S55. These residues form direct contacts with the ligand and contribute to the overall stability of the complex. In sum, the docking investigations highlight a specific binding region on LysB_FL_ that interacts effectively with (GlcNAc)_5_, thus shedding light on the putative binding mechanism of LysB_FL_.

### LysB_FL_ is efficiently taken-up by murine alveolar-like macrophages (mAMs) and triggers a mild pro-inflammatory state

Considering that AMs are the natural niche for *M. tuberculosis*, and essentially the are the first line of defense against this pathogen^20^, we sought to assess whether LysB_FL_ can be internalized by AMs and whether the protein modulates the AMS polarization state. As a cellular model, we adopted a murine cell line, namely Max Plank Institute-2 (MPI-2), that closely reproduces the AMs-like phenotype^21,22^. First, we assessed the internalization of LysB_FL_ in MPI-2 cells, thereafter referred to as mAMs, by high-resolution confocal microscopy. We observed the progressive accumulation of LysB_FL_ over time and the lack of accumulation of the fluorescent tag alone (sfGFP) (Figure 3A**, 3B** and **S2B**). Importantly, the uptake of the recombinant protein did not induce any significant alteration in cell viability, even after 24 hours incubation at the highest concentration tested (15 nmol) (**Figure S2A**). We also investigated whether the internalization of LysB_FL_ promote alterations of mAM polarization by assessing gene expression of several pro-and anti-inflammatory markers. We incubated the cells with the highest concentration of recombinant protein for 24 hours to ensure a significant intracellular accumulation of LysB_FL_. Overall, the recombinant protein inhibited the expression of anti-inflammatory markers (*Stat6*, *CD20* and *Tgfβ*) and significantly increased pro-inflammatory genes (*Stat1*, Il-1*ý, Il-6,* Il-1*α* and *Tnfα*) (Figure 3C).

**Figure 3.**
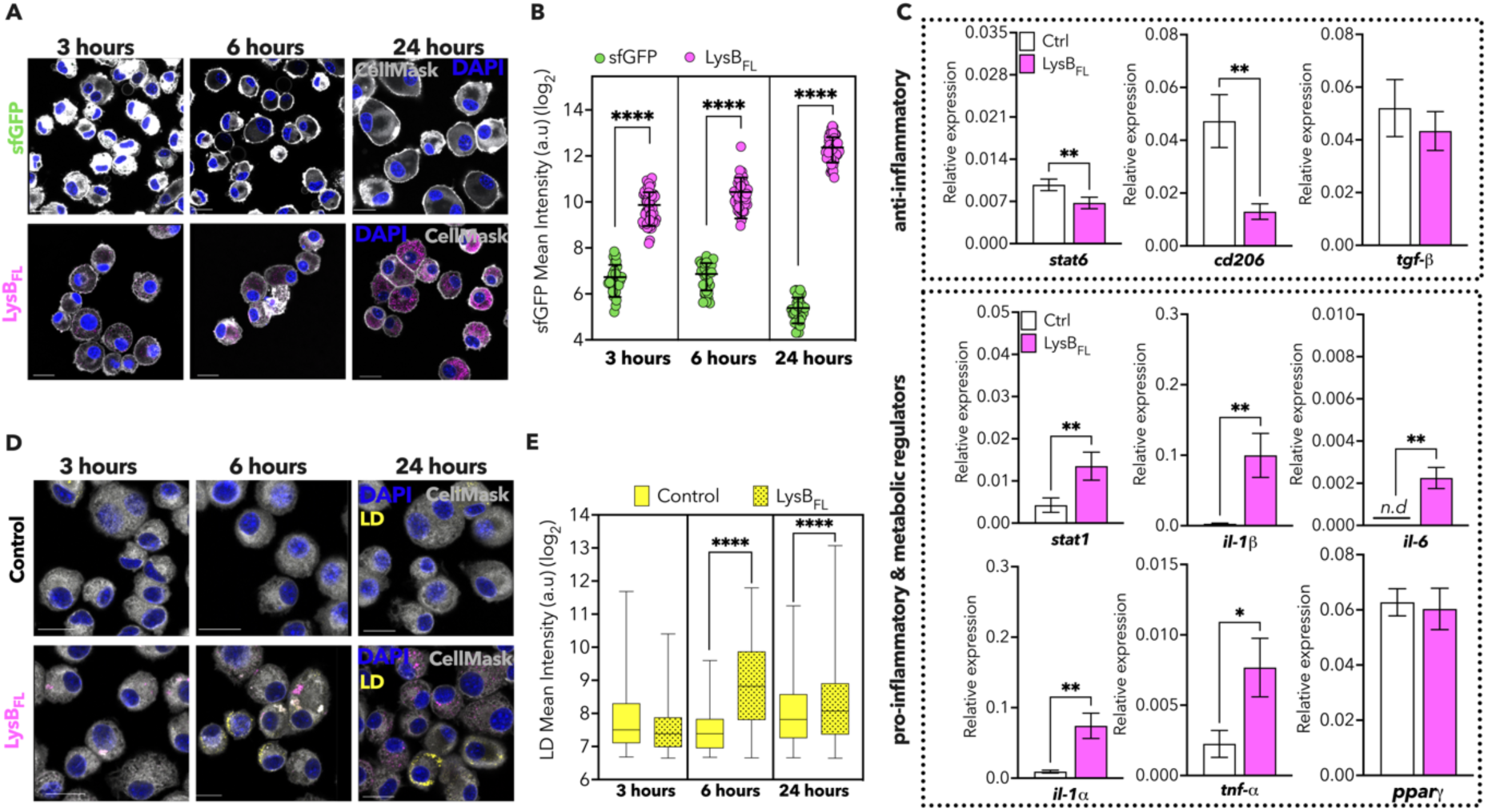
LysB_FL_ might promote a pro-inflammatory polarization in AMs-like cells. (A) Representative images of sfGFP (upper panel) and LysB_FL_ (lower panel) uptake kinetics in murine AMs-like cells. AMs-like cells (gray), sfGFP (green), LysB_FL_ (magenta), and DAPI (blue) are merged. Scale bar 20 μm. (B) Quantification of sfGFP (green) and LysB_FL_ (magenta) cellular entry over time in AMs-like cells. Black lines indicate mean value ± SD (30 > n <57). Asterisks denote significant differences by One-way ANOVA followed by Turkey’s multiple comparison test: ****p< 0.0001 (C) qRT-PCR analysis comparing the expression of selected pro-and anti-inflammatory cytokines in AMs-like cells either treated or not with 15 nmol of LysB_FL_ (magenta) for 24 hours. AMs-like cells in optimal condition were used as control (white). Transcripts are normalized to total RNA and gapdh expression. Data are shown from 4 independent experiments. Mean value ± SD are plotted (n =4). Asterisks denote significant differences by One-way ANOVA followed by Turkey’s multiple comparison tests: *p = 0.01, **p = 0.002, ***p =0.0009. (D) Representative images of lipidic droplet (LD) formation in murine AMs-like cells treated (lower panel) or not (lower panel) with 15 nmol of LysB_FL_. AMs-like cells (gray), LysB_FL_ (magenta), and DAPI (blue), LD (Nile Red, yellow) are merged. Scale bar 20 μm. (E) Box plot quantifying LD formation in AMs-like cells treated (lower panel) or not (upper panel) with 15 nmol of LysB_FL_ over time in AMs-like cells. Black lines indicate mean value ± SD (37 > n < 2050). Asterisks denote significant differences by One-way ANOVA followed by Turkey’s multiple comparison test: ****p < 0.0001.

To further confirm the effect of LysB_FL_ in promoting a pro-inflammatory polarization of AMs, we assessed the intracellular accumulation of lipid droplets (LDs), which is associated to a pro-inflammatory phenotype^25,26^. We incubated mAMs with 15 nmol of either LysB_FL_ or sfGFP for 3, 6, and 24 hours and stained intracellular LDs using Nile Red dye. We observed a limited formation of LDs after 24 hours in steady state (untreated) mAMs, while incubation with LysB_FL_ triggers a large number of cytoplasmic LDs, which peaked 6 hours after LysB_FL_ treatment (Figure 3D**, 3E** and **S2C**). We also observed a change in mAMs shape when we compared cells treated with either sfGFP or LysB_FL_ (**Figure S2D**), possibly suggesting a diverse degree of either sfGFP or LysB_FL_ variation in mAMs phenotype. Overall, our data suggests that LysB_FL_ reshapes mAMs phenotype promoting the polarization towards a pro-inflammatory phenotype.

### LysB_FL_ might help mAMs in controlling *M. tuberculosis* infection

Accounting for LysB_FL_ affinity for *M. tuberculosis* OM and its ability to naturally permeate within mAMs inducing a partial pro-inflammatory polarization, we hypothesized that LysB_FL_ could aid mAMs to better respond to *M. tuberculosis* infection. First, we investigated whether the uptake of the recombinant protein in *M. tuberculosis* infected mAM is affecting cell viability at either 6 or 24 hours of incubation, showing only a significant reduction in infected and LysB_FL_-treated mAMs viability at the latest analyzed time point (**Figure S3A**) Parallelly, we carried out gene expression analysis in mAMs either infected or not, and then treated with 15 nmol LysB_FL_ overnight. Interestingly, we observed that infected mAMs treated with LysB_FL_ strongly expressed pro-inflammatory genes, such as Il-1α, Il-1ϐ, Il-6, cytokines important for the control of *M. tuberculosis* spreading. Moreover, exposure to LysB_FL_ also increased the expression of Tgf-ϐ, which is known to play a crucial role in regulating *M. tuberculosis* infection, in preventing hyperinflammation and autoimmunity_27_. In this regard, and although still in its infancy, there is emerging evidence that while bacteriophages could explicit their antimicrobial functions, they parallelly communicate with the cells of the immune system (*i.e.,* macrophages and lymphocytes) to contribute to the maintenance of immune homeostasis^28,29^. Hence, pointing out that such event might depend on the putative LysB_FL_ immune modulatory effect^6,7^.

Next, we used high-resolution confocal microscopy to better understand the role of LysB_FL_ during *M. tuberculosis* infection (Figure 4B and **S3B**). By setting a multiparameter single-cell image analysis, we discriminated between mAMs hosting intracellular *M. tuberculosis*, namely infected, and cells that lacked the presence of the bacilli but were exposed to it, here referred to as bystanders. We used LysB_FL_ fluorescence as a proxy to quantify the intracellular accumulation of the recombinant protein in infected and bystander cells. Although we observed a comparable distribution of LysB_FL_ fluorescence in the two subpopulations (**Figure S3C**), infected mAMs were characterized by a lower, but significant, intracellular accumulation of LysB_FL_ (**Figure S3D**). Despite the reduced accumulation inside infected AMs, treatment with the recombinant protein promoted a slight reduction in the percentage of infected cells compared to the untreated control (Figure 4C). To examine whether the reduction of infection rate is a direct consequence of LysB_FL_ antimicrobial activity, or more broadly due to the activation of the cells, we quantified the number of *M. tuberculosis* bacilli in AMs treated with LysB_FL_ by confocal microscopy. We quantified a significant reduction in the number of intracellular pathogens after treatment with LysB_FL_ (p<0.01) (Figure 4D). Furthermore, image analyses revealed the presence of sfGFP fluorescence around *M. tuberculosis* bacilli inside infected cells. This indicates the association between the recombinant protein and the pathogen (Figure 4D, **4F** and **S3E)**.

**Figure 4.**
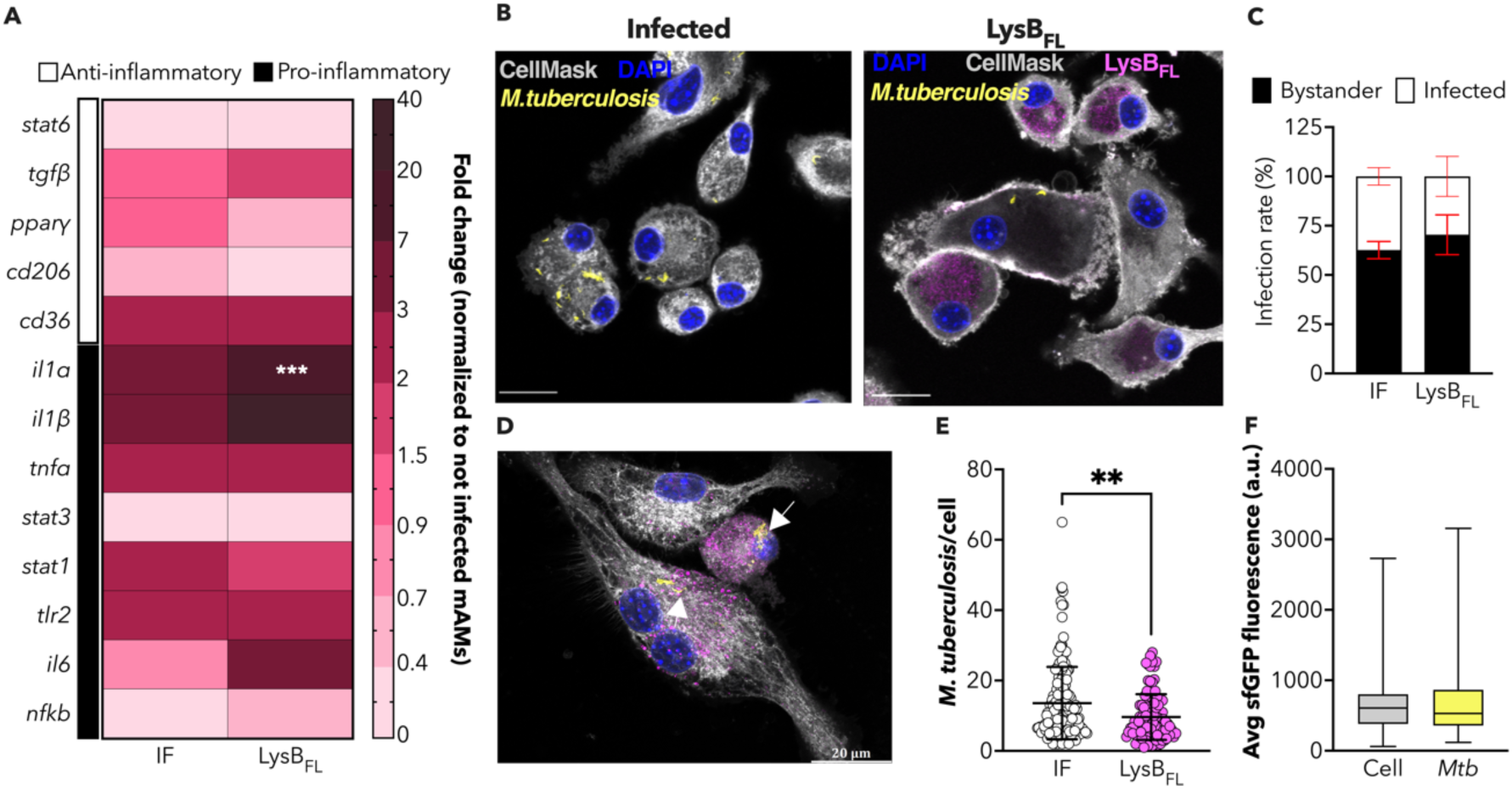
LysB_FL_ might help mAMs to better control M. tuberculosis infection. (A) Heat map showing relative gene expression of selected anti-(white) and pro-inflammatory (black) markers in mAMs infected with M. tuberculosis and then treated or not with 15 nmol of LysB_FL._ Transcripts are normalized to total RNA and gapdh expression. Data are shown from 4 independent experiments. Asterisks denote significant differences with respect to the not-infected control by Two-way ANOVA followed by Dunnett’s multiple comparison test: ****p< 0.0001. (B) Representative images of mAMs infected with M. tuberculosis and treated (left panel) or not (right panel) with 15 nmol of LysB_FL_. AMs-like cells (gray), LysB_FL_ (magenta), M. tuberculosis (yellow), and DAPI (blue) are merged. Scale bar 20 μm. (C) Infection rate expressed as the percentage of infected mAMs with M. tuberculosis four hours post-infection. Infected mAMs are indicated in white, and bystander cells in black. Data are shown from four independent experiments. Red lines indicate Mean value± SD (n=4) (D) Maximum projection of mAMs infected with M. tuberculosis and treated with 15 nmol of LysB_FL_. White arrows indicate bacilli and LysB_FL_ proximity. AMs-like cells (gray), LysB_FL_ (magenta), M. tuberculosis (yellow), and DAPI (blue) are merged. Scale bar 20 μm. (E) Number of intracellular M. tuberculosis per single mAMs detected in infected and not treated mAMs (white) and in infected mAMs treated with 15 nmol LysB_FL_ (magenta). Data are shown from four independent experiments. Asterisks denote significant differences by Mann Whitney test: **p=0.0012. (F) Box plot comparing sfGFP fluorescence average detected within the cytoplasm of AMs cells infected (light gray) and treated with 15 nmol of LysB_FL_ and sfGFP fluorescence average measured over M. tuberculosis area. Data are shown from four independent experiments.

Overall, our findings suggest that LysB_FL_ might be actively limiting the spread of the infection by modulating mAMs polarization and by directly promoting *M. tuberculosis* intracellular clearance as it binds to the engulfed bacilli and induces their elimination. Our results confirm the concerted antimicrobial action of LysB_FL_, whicle mAMs might contribute to better control of *M. tuberculosis* dissemination and infection progression.

## Discussion

One of the mycobacteria’s distinctive characteristics is their complex and lipid-rich cellular envelope, commonly known as mycomebrane^30^. In particular, the mycomembrane is highly enriched in mycolic acids, long-chain fatty acids, which are esterified to the underlying arabinogalactan layer, with intercalating glycol-and phospholipids and fatty acid^30^. Hence, constitutes a near-impermeable structure, which heavily contributes to mycobacterial intrinsic resistance to most of the commonly used antimicrobials^23,30^. Additionally, the presence of this thick waxy layer further complicated mycobacteriophages, bacterial viruses selectively parasitizing mycobacteria, lysis^23^. Generally, bacteriophages encode within their genome only one peptidoglycan-degrading enzyme, namely endolysin A (LysA), which is essential in allowing bacteriophage extracellular propagation at the end of their lytic cycle^31,32^. Although the molecular mechanisms triggering peptidoglycan endolysins-mediated lysis are still poorly elucidated, the current model hypothesized that LysA starts to inertially accumulate at the inner membrane together with holins. This can lead to a possible drop in the bacterial proton motive force, a process triggering holin to create micron-scale holes at the membrane level. Subsequently to the formation of these micro-fractures, LysA can finally reach the bacterial cell wall and start degrading the peptidoglycan^33,34^. Yet, in mycobacteria, the release of newly assembled phages destroys the mycolic acid layer^12,16^. This portrays an additional degree of complexity to viral particle extrusion from the host. To overcome the complexity of the mycomembrane, mycobacteriophages release two different endolysins within the cytoplasm of their host, namely LysA and the mycolylarabinogalactan esterase endolysin B, LysB^12,13^. While LysA acts hydrolyzing the peptidoglycan layer, LysB cuts the ester link between the arabinogalactan and the mycolic acid disrupting mycobacterial integrity^32^. The concerted degrading action of both LysA and LysB finally leads to mycobacterial lysis, which occurs following the loss of communication between the bacillus cell wall the outer membrane^32^. Therefore, the endolysin-mediated lytic activity towards host bacteria, combined with the pressing need to counterbalance the exponential rise of antibiotic-resistant pathogens, are posing these lytic-derived enzymes as new potential antibacterial _agents_^18,23,32^.

In this study, we combined computational chemistry, biochemistry, and high-resolution confocal imaging to characterize Ms6-encoded LysB_FL_ mycomembrane-binding properties both qualitatively and quantitatively, while shedding light on its ability to aid mAMs in controlling *M. tuberculosis* infection.

Initially, we generated a fluorescent LysB_FL_ to confirm binding to the OM of both *M. smegmatis* and *M. tuberculosis*^12,13,16,24^. However, our observations revealed a distinct binding specificity of LysB_FL_ towards *M. tuberculosis*, possibly due to an intrinsic higher affinity towards slow-growing pathogenic rather than fast-growing non-pathogenic mycobacteria. While we observed that LysB_FL_ could bind to around 40% of the *M. smegmatis* population, almost the totality of *M. tuberculosis* suspension indeed proved to be positive for LysB_FL_. Although *M. smegmatis* is often used as a model organism for pathogenic mycobacteria, it is actually distantly related to them. On the structural level, the different specificity of binding could be due to the presence of glycopeptidolipids in the mycomembrane of *M. smegmatis* and hydrophobic lipids like phthiocerol dimycocerosates (DIM/PDIM) on *M. tuberculosis*^23,35,36^. This diverse composition could potentially modulate the overall affinity of LysB_FL_.

To date, the two mostly investigated mycobacteriophage lysins are the Ms6-LysB_FL_ and the D29-LysB^18,32^. Whereas recombinant D29-LysB appears to induce mycobacterial lysis and to disrupt biofilm^32^, Ms6-LysB_FL_ only showed a growth-inhibitory effect when co-administered with a surfactant or with membrane-destabilizing cationic peptides^12,16,23,37^. Contrary to this previous evidence, we demonstrated that Ms6 LysB_FL_ can significantly affect the growth of planktonic cultures of *M. tuberculosis* even in the presence of low concentration of polysorbate 80 (0.05%). This possibly indicates that even a minimal destabilization of the external capsule, which might restrict LysB_FL_ binding^23^, could allow LysB_FL_ penetration within the mycolic acid layer. This would promote cell wall hydrolysis and the inhibition of *M. tuberculosis* growth, thus further supporting LysB_FL_ antimycobacterial effect. In this regard, because LysB_FL_ acts on non-genomic encoded targets (*i.e.,* mycolic acid), diversely from the clinically used antimicrobials, the rise of resistance to their lytic activity is rare and could be exploited to generate innovative and specific antimicrobial therapies^18,38–40^. On the other hand, one of the main limitations to the use of phage and/or phage-derived proteins for therapy stands in the possible immune response that the host could develop in response to exogenous proteins^41,42^. Although the administration of isolated and bioengineered endolysins ensured rapid effects on Gram-positive bacteria that resulted in various clinical trials^43,44^, these proteins have also been associated to the secretion of inflammatory cytokines and neutralizing antibodies, which significantly reduced the endolysin half-life and the loss of its enzymatic lytic activity^43^. Consistent with these findings, mAMs treated with LysB_FL_ tend to polarize towards a pro-inflammatory phenotype. Indeed, we not only detected an increase in the production of cytokines such as *Il-1*α, *Il-1ϐ,* and *Tnf-*α but also an enhanced intracellular accumulation of lipid droplets, molecular indicators of macrophage activation^45^. Moreover, while we could detect LysB_FL_ internalization, which is heavily dispersed within the cytoplasm of mAMs, we could not quantify any intracellular sfGFP fluorescence. In this regard, although endolysins seem to conserve their lytic activity intracellularly when used in *in vivo* systems^46–48^, little is known about their ability, either intrinsic or conferred by molecular engineering, to penetrate the eukaryotic membrane. Hence, our data could potentially suggest that LysB_FL_ entry might undergo selective and active endocytosis. Yet, the molecular mechanism(s) potentially driving such a process remains unrevealed. Overall, our data indicate that LysB_FL_ has an intrinsic immune-modulatory effect, which induced mAMs towards a pro-inflammatory phenotype. However, LysB_FL_ appears to induce the expression of anti-inflammatory factors that might result in only a mild mAMs activation. The high affinity displayed by LysB_FL_’s towards *M. tuberculosis*, and the absence of a significant cytotoxic on mAMs, prompted us to investigate whether LysB_FL_ could also help mAMs in controlling and even eradicating the intracellular bacterial., We showed, transcriptomics analysis, that LysB_FL_-treated and infected mAMs exhibited a higher pro-inflammatory profile than control. However, mAMs could potentially avoid a hyperinflammatory state, which might favor *M. tuberculosis* dissemination and infection progression^49^, by parallelly secreting anti-inflammatory factors, such as TGF-*ϐ*. We then used high-resolution confocal microscopy to test whether LysB_FL_ retains its lytic mycobacterial activity even within mAMs intracellular environment, and to infer whether LysB_FL_ promotes pathogen clearance. Although we could not detect any significant reduction in the percentage of infected cells upon treatment with LysB_FL_, we observed a significant reduction in the number of intracellular bacilli. Hence, we can speculate that LysB_FL_ can potentially retain its lytic activity, which promotes *M. tuberculosis* clearance.

Finally, our computational atomistic model of the LysB_FL_-(GlcNAc)_5_ complex yields crucial insights into the binding mode of LysB. Notably, LysB exhibits a shallow negative patch, pinpointed as a potential binding site through blind docking simulations. Within this region, various residues on the surface of LysB are characterized for their involvement in the interaction with the GlcNAc carbohydrate. This detailed analysis provides a comprehensive understanding of the specific molecular interactions and highlights key amino acid residues for the crucial binding affinity and specificity in the LysB-GLcNAc interactions.

In conclusion, LysB_FL_ affinity for its host both *in vitro* and in mAMs corroborates its intrinsic ability to induce *M. tuberculosis* lysis while possibly helping mAMs in better controlling the infection. *M. tuberculosis* continues to take a heavy toll on global public health and innovative strategies are urgently needed to limit the spreading of this drug-recalcitrant infection^50^. Thus, the lytic antimycobacterial effect exerted with high selectivity by LysB_FL_ might represent a promising therapeutic tool to treat TB. Importantly, this approach could easily be tuned, by selecting other LysBs, and implemented to create a platform of therapies to treat infections caused by other mycobacteria.

## Acknowledgments

We thank the OMICs Mass spectrometry platform for their proteomic analysis. The authors also acknowledge the scientific and technical assistance of Dr Alessandra Fasciani and Dr Chiara Cordiglieri of the INGM Imaging Facility, and Dr Mariacristina Crosti of the Sorting Facility at INGM. We also thank Prof. Marina Freuderberg (BIOOS Centre for Biological Signalling Studies, University of Freiburg) for kindly providing MPI-2 cells. We also would like to thank Dr Giacomo Butta (INGM, Virology Laboratory) and Chiara Zara (Milteniy) for their help with the Tyto® sorter. This work was funded by grants to L.R from ERC-2019-STG (grant number 850936), the Fondazione Cariplo (grant number 2019-4278), and the Italian Ministry for University and Research (through the PRIN program, grant number 20205B2HZE).

## Author contributions

Conceptualization L.R, A.G and E.S. Methodology A.G, E.S, G.M, A.S, M.G, R.M, S.R and L.R. Data curation A.G, G.M, E.S, A.S, M.G, R.M, B.A, S.M and L.R. Validation A.G, G.M, E.S, A.S, B.A, S.M, I.M, M.G, and L.C. Formal Analysis A.G, A.S, B.A, S.M, M.G, R.M, G.M and E.S. Investigation A.G, B.A, A.S, S.M, M.G, I.M, G.M. Resources A.G, A.S, G.M, E.S, B.A, S.M, M.G, R.M, I.M, L.C, L.R. Writing – Original draft A.G, E.S, A.S, and L.R; Writing – Review & Editing A.G, S.A., B.A, G.M, E.S, S.M, L.C, I.M, M.G, V.D.M., R.M,S.R, L.R. Visualization A.G, A.S, G.M, E.S, V.D.M., M.G, R.M and L.R. Supervision A.G, E.S, G.M, and L.R. Project administration A.G, E.S and L.R. Funding acquisition L.R.

## METHODS

### RESOURCE AVAILABILITY

#### Lead contact

Further information and requests for resources and reagents should be directed to and will be fulfilled by the Lead Contact, Loris Rizzello

#### Materials Availability

Plasmids and bacterial strains generated in this study are available from the Lead Contact with a completed Materials Transfer Agreement

### EXPERIMENTAL MODEL AND SUBJECT DETAILS

#### Bacterial strains and growth conditions

All mycobacterial strains were cultured at 37°C in Middlebrook 7H9 broth (Sigma) supplemented with 10% OADC (Fisher Scientific), 0.5% glycerol (Sigma), and 0.05% Tween-80 (Sigma). Middlebrook 7H10 agar (Sigma) has been enriched with 10% OADC and 0.5% glycerol. Exponentially growing cultures were started from a single colony, aliquots were enriched with 15% glycerol, stored at −80°C, and used once to start primary cultures.

*M. smegmatis* and *M. tuberculosis* were grown in Middlebrook 7H9 broth at 37°C in shaking condition, 150 and 50 rpm, respectively, to a mid-log phase (OD_600_ 0.5-0.8) before any experiments, unless specified otherwise. The selective antibiotic was only added for strains carrying chromosomal integrative vectors in the primary culture and removed in the secondary culture used for the final experiments.

Constitutive reporter strains used in the following study were generated using the integrative vector pTdTomato-L5 (Addgene #140994). The chromosomal integration was done via the L5 phage site *attB*^51^. Both mycobacterial species were transformed with the construct by electroporation (2 500 V, 25 μF, 1000 Ω, 2 mm path), and plated on Middlebrook 7H10 agar containing Streptomycin (Sigma) at a concentration of 30 μg/mL. *M. smegmatis::tdTomato* and *M. tubercolisis::tdTomato* were named AGS1 and AGT1, respectively.

#### Strains construction

The plasmid encoding sfGFP was generated by ordering its synthesis and cloning into the pET23a(+) expression vector (Vectorbuilder). In the final construct, sfGFP was also C-tagged with a 6xHist tag and its expression was under the control of a T7 inducible promoter.

The plasmid encoding Ms6 LysB was generated by ordering the synthesis of Ms6 *lysB* ORF and cloning it into the pET23a(+) expression vector (GeneScript). Additionally, the Ms6 *lysB* ORF was fused at the C-terminal with sfGFP. An oligopeptide linker was added between the protein and the fluorescent reporter to not affect the folding of the protein. Between the Ms6 LysB and the sfGFP, an *EcoRI* restriction site was added. Finally, the LysB C-fused with sfGFP was cloned in-frame into the pET23a(+) vector between the restriction site *NheI* and *NotI*. In the final construct, Ms6 LysB-sfGFP (LysB_FL_) was also C-tagged with a 6xHist tag and its expression was under the control of a T7 inducible promoter.

#### Protein expression and purification

For the expression of the recombinant sfGFP and LysB_FL_ *E. coli* BL21(DE3) strain (NEB) were used. 50 μl of BL21(DE3) were transformed with a maximum of 50 ng of pET23a_LysB_FL_ or pET23a_sfGFP, respectively. The transformed cells were then plated on LB agar with the selection (100 μg/mL ampicillin) and incubated overnight at 37°C.

One transformed colony was then used to prepare a 20 mL overnight liquid culture to which the section agent (100 μg/mL ampicillin) was added. To induce the expression of either sfGFP or LysB_FL,_ 1L of LB culture with a starting OD_600_ of 0.02 was prepared. Each hour, the OD_600_ was measured, and 1 mL of culture was stored for downstream analysis. When the OD_600_ reached values of 0.6, the over-expression of the recombinant proteins was induced by adding a final concentration of 100 µM of IPTG. The induction of the recombinant proteins was then continued for 3 hours. Then, bacterial cells were harvested by centrifugation at 10 000 *xg* for 10 minutes, and the pellet was stored at −20°c. The collected pellets were lysated by chemical lysis using B-PER^TM^ Bacterial Protein Extraction buffer (ThermoFisher). For each gram of pellet, 2 ml of B-PER^TM^ Bacterial Protein Extraction buffer was used. Finally, the pellets were incubated for 15 minutes at room temperature. Cell debris were removed by centrifugation at 20 000 *xg* at 4°C for 30 minutes, and the supernatant containing the recombinant proteins was collected. SfGFP and LysB_FL_ purification was performed using a 3 mL HistPur^TM^ Cobalt Purification Kit (ThermoFisher), according to the manufacturer’s instructions.

#### sfGFP and LysBFL concentration

The purified proteins were stored in PBS at −20°C. sfGFP and LysB_FL_ were desalted performing 3 washes in PBS using Pierce^TM^ Protein Concentrator PES, 30 K MWCO (ThermoFisher). Pierce^TM^ Protein Concentrator PES, 30 K MWCO (ThermoFisher) were equally used to concentrate the recombinant proteins to a final concentration of at least 5 mg/mL. Protein concentration was determined by the Bradford (Sigma) method using 2 mg/mL BSA as a standard (ThermoFisher).

#### Coomassie staining

10% separating gel was prepared with 1.5 mL dH_2_O, 3.3 mL of 30% acrylamide-bisacrylamide (Sigma), 5.0 mL of 2x Separating buffer (2.5 M Tris, pH 8.8, 10% SDS), 100 μl of 10% APS (Sigma) and 10 μl of TEMED (Sigma). 4% staking gel was prepared with 1.4 mL of dH_2_O, 2.0 mL of 30% acrylamide-bisacrylamide (Sigma), 2.0 mL of 2x Separating buffer (0.5 M Tris, pH 6.8, 10% SDS), 40 μl of 10% APS (Sigma) and 4 μl of TEMED (Sigma). Before loading 6x loading buffer (ThermoFisher) was added to the samples and then the samples were incubated for 10 minutes at RT prior to their loading. The gels were run for c.a 2 hours with a constant electric current of 100V in 1x NuPAGE running buffer (ThermoFisher). The SDS-PAGE was then stained with Coomassie. The gel was first washed with water and then incubated with c.a 10 mL of InstantBlue^TM^ Ultrafast Protein Stain (Sigma) for at least 1 hour.

#### Western blot

Western blot assays were carried out either on whole cell extract after induction or on the purified sfGFP and LysB_FL_. The samples were run on a 10% SDS-PAGE for c.a 2 hours at 100V. Proteins were then transferred into a PVDF membrane (Sigma) by wet transfer at 100V for 1 hour. The membrane was blocked with 5% Milk (Sigma) prepared in 1x Tween20 TBS (ThermoFisher) for at least 1 hour, incubated with primary antibody overnight at 4°C, washed three times with TBS-T for 5 minutes, incubated with peroxidase-conjugated secondary antibody in 5% Milk TBS-T for at least 1 hour at RT, and washed three times with TBS-T for 5 minutes. Proteins were detected using the Pierce^TM^ ECL Western Blotting Substrate (ThermoFisher) and InvitrogenTM iBright_TM_ 1500 series. Primary and secondary antibodies were diluted in TBS-T and 5% milk as follows: anti-GFP (SantaCruz Biotechnology) 1:200, anti-6xHist (SantaCruz Biotechnology) 1:100, HRP-conjugated anti-mouse (SantaCruz Biotechnology) 1:2000.

#### LysB Mass Spectrometry

We conducted qualitative Mass Spectrometry analysis to confirm LysBFL (ID uniport Q9FZR9) protein sequence. Thus, two samples of purified LysB_FL_ were concentrated by Trichloroacetic acid (TCA)/acetone purification. All the steps of the TCA/Acetone precipitation were performed at 4°C if not otherwise specified. To each sample, 1 volume of cold 40% TCA (Sigma) was added, followed by 30 minutes of incubation, and then centrifuged for 15 minutes at 14 000 *xg*. The pellet was then washed with 1 volume of cooled acetone (Sigma), incubated for 10 minutes, and centrifugated for 10 minutes at 14 000 *xg*. After the final washing step, the supernatant was discarded, and the sample dried. The precipitated samples were stored at 4°C before Mass Spectrometry.

After TCS/acetone precipitation the samples were resuspended in 50 mM ammonium bicarbonate (Ambic) and then 20 μl of each sample was then digested with trypsin. Digested samples were analyzed using Dionex Ultimate^TM^ 3000 nano-LC system (Sunnyvale) connected to Orbitrap Fusion^TM^ Tribrid^TM^ Mass Spectrometer (Thermo Scientific) equipped with nano electrospray ion source. Peptide mixtures were pre-concentrated into Acclaim PepMap 100-100 μm x 2 cm C18 (Thermo Scientific) and separated on EASY-Spray column ES802A, 25 x 75 μm ID packed with Thermo Scientific Acclaim PepMap RSLC C18, 3 μm, 100 Å using mobile phase A (0.1% formic acid in water) and mobile phase B (0.1% formic acid in acetonitrile 20/80, v/v) at a flow rate of 0.300 μL/min. The temperature was set to 35°C and the sample were injected in duplicate. MS spectra were collected over an m/z range of 375 – 1500 Da at 120 000 resolutions, operating in the data dependent mode, cycle time 3 sec between master’s scans. HCD was performed with collision energy set at 35 eV. Polarity: positive.

#### Plate binding assay

*M. tuberculosis* and *M. smegmatis* constitutively expressing tdTomato were cultured at 37°C in complete Middlebrook 7H9 broth in shaking conditions (50 and 150 rpm, respectively) until the mid-log exponential phase (OD_600_ 0.5-0.8). At this point, the mycobacterial suspension was centrifuged at 3 900 rpm for 15 minutes at room temperature. Then the pellet was resuspended in filtered PBS, and bacteria were diluted to obtain serial dilution with the following OD_600_ 7, 6, 5, 4, 3, 2, 1, 0.5. The samples were then centrifuged for 5 minutes at 15 000 rpm, and the pellets were resuspended with 50 μl of either sterile LysB_FL_ or sfGFP proteins with a final concentration of 15nmol. Next, the samples were incubated for 3 hours at room temperature. After 3 hours, the treated samples were centrifugated at 15 000 rpm for 5 minutes, and the unbound protein was discarded. To remove eventual unspecifically bound protein, the samples were washed twice with 50 μl of PBS 0.05%Tween80. Finally, the pellets were resuspended in 50 μl of PBS 0.05%Tween80 and moved into a 96 black multi-well plate (Thermo Scientific), and the assay was then analyzed by reading sfGFP and tdTomato fluorescence with a plate reader (TECAN, Spark).

Parallelly, calibration curves for the bacteria, LysB_FL_ and sfGFP were prepared and used to analyze the data.

#### Microscale Thermophoresis (MST)

*M. smegmatis* mc^2^155 was cultured at 37°C in complete Middlebrook 7H9 broth in shaking conditions (150 rpm) until mid-log exponential phase (OD_600_ 0.5-0.8). At this point, the mycobacterial suspension was centrifuged at 3 900 rpm for 15 minutes at room temperature. Then the pellet was resuspended in filtered PBS, and bacteria were diluted to obtain serial dilution with the following OD_600_ 7, 6, 5, 4, 3, 2, 1, 0.5. The samples were then centrifuged for 5 minutes at 15 000 rpm, and the pellets were resuspended with 50 μl of either sterile LysB_FL_ or sfGFP proteins with a final concentration of 1.25 nmol. Next, the samples were incubated for 3 hours at room temperature. After 3 hours, around 5 μl of the samples were loaded in Monolith^TM^ Capillaries and MST analysis was carried out on the Instrument (Nanotemper).

#### Surface Plasmon Resonance (SPR)

SPR measurements were performed at 20°C using a Biacore X100 instrument (Cytiva). *M. smegmatis* mc^2^155 samples used for SPR experiments were cultured at 37°C in complete Middlebrook 7H9 broth in shaking conditions (200 rpm) until OD600 0.3 (#1) and OD600 0.8 (#2). LysBFL protein was immobilized onto a CM5 sensor chip functionalized with an anti-His antibody attached by amine-coupling (following manufacturer instructions, Cytiva), reaching a target capture level of 76.1 Resonance Units (RUs). Once the immobilization procedure was completed, non-specifically bound proteins were removed by washing with HBS-N buffer (Cytiva) until the resonance signal became nearly constant. The reference flow cell was used as a control surface for refractive index change and non-specific binding. The analysis has been performed by manual injection of the two mycobacterial samples by means of Biacore X-100 instrument (Cytiva) and HBS-N buffer as mobile phase.

#### Flow Cytometry assay

*M. smegmatis* mc^2^155 was cultured at 37°C in complete Middlebrook 7H9 broth in shaking conditions (150 rpm) until mid-log exponential phase (OD_600_ 0.5-0.8). At this point, the mycobacterial suspension was centrifuged at 3 900 rpm for 15 minutes at room temperature. Then the pellet was resuspended in filtered PBS 0.05% Tween80, and bacteria were diluted to obtain serial dilution with the following OD_600_ 7, 6, 5, 4, 3, 2, 1, 0.5. The samples were then centrifuged for 5 minutes at 15 000 rpm, and the pellets were resuspended with 50 μl of either sterile LysB_FL_ or sfGFP proteins with a final concentration of 15 nmol. Next, the samples were incubated for 3 hours at room temperature. After 3 hours, to remove eventual unspecifically bound protein, the samples were washed twice with 50 μl of PBS 0.05%Tween80. Finally, the pellets were resuspended in 300 μl of PBS 0.05%Tween80 and analysed with BD FACSCanto^TM^ flow cytometer. Physical cellular parameter (*i.e.,* FSC-A and SSC-A) were set up using commercially available Megamix beads (Cosmo Bio Co, LTD) according to the manufacturer’s instructions.

*M. tuberculosis* was cultured at 37°C in complete Middlebrook 7H9 broth in shaking conditions (50 rpm) until mid-log exponential phase (OD_600_ 0.5-0.8). At this point, the mycobacterial suspension was centrifuged at 3 900 rpm for 15 minutes at room temperature. Then the pellet was resuspended in filtered PBS 0.05% Tween80, and bacteria were diluted to obtain serial dilution with the following OD_600_ 7, 6, 5, 4, 3, 2, 1, 0.5. The samples were then centrifuged for 5 minutes at 15 000 rpm, and the pellets were resuspended with 50 μl of either sterile LysB_FL_ or sfGFP proteins with a final concentration of 15 nmol. Next, the samples were incubated for 3 hours at room temperature. After 3 hours, to remove eventual unspecifically bound protein, the samples were washed twice with 50 μl of PBS 0.05%Tween80. Finally, the pellets were resuspended in 500 μl of MaCSQuant® Tyto® running buffer and analysed with MACSQuant® Tyto® Cell Sorter (Miltenyi Biotec).

#### Molecular dynamics and docking calculations

LysB_FL_ structure has been modelled using AlphaFold2 version 2.3.2, using the follow fasta sequence: *M(1)*RIDGQYVGLGPGDRSDEIRKIKAFMRRKFSYAATLADTEFYDEAMTAVVAEMQSRYNTAGQLRDGLYIPGIINAETKYVMGYLSRPVIDTRPVLFTVCGTGVPWWVGPDADTARAVEDQYLWQPIGYPAAPFPMGRSITAGITEAHNQANRWRERIETHGTALAGYSQGAVVLSELWMNHIAPEDGSLRWMKPHVRKAVTWGNPNRELGHVWADHGGSPMAPSNTQGVSSNGMRNTPDWWRDYAHQGDLYACTEPGDTQEVRNAIWQIVRDLDLFTGPDSLLAQVIELAQAPLPETIAITRAILDAGMFFAKRTGPHVDYNPQPAIDYLRT3*30G331 SGSGS337M338*SKGEELFTGVVPILVELDGDVNGHKFSVRGEGEGDATNGKLTLKFICTTGKLPVPWPTLVTTLTYGVQCFSRYPDHMKRHDFFKSAMPEGYVQERTISFKDDGTYKTRAEVKFEGDTLVNRIELKGIDFKEDGNILGHKLEYNFNSHNVYITADKQKNGIKANFKIRHNVEDGSVQLADHYQQNTPIGDGPVLLPDNHYLST QSVLSKDPNEKRDHMVLLEFVTAAGITHGMDELY*K576* where 1-330 represents LysB, 331-337 the linker and 338-576 sfGFP protein sequences respectively. AlphaFold2 calculations were performed on LysB and LysB_FL_ using the monomer_casp14 model to evaluate the fold of the LysB in presence or absence of sfGFP. The best ranked model from LysB_FL_ was used in the following molecular dynamics simulations and docking calculations.

Molecular dynamics simulations have been performed using GROMACS-2020.2. The Amber14 force field was used with TIP3P waters model. Adding a suitable number of counter-ions neutralized the overall system. After energy minimization, four consecutive equilibration steps were then performed:1) 100ps in the NVT ensemble at 100K with the protein backbone restrained (k=1000 kJ/mol nm2), 2) 100ps in the NVT ensemble at 200K with the protein backbone similarly restrained, 3) 100ps in the NVT ensemble at 300K with the protein backbone restrained, and 4) 1000-ps in NPT ensemble at 300K with no restraints. For atoms less than 1.1nm apart, electrostatic forces were calculated directly; for atoms further, apart electrostatics were calculated using the Particle Mesh Ewald. Van der Waals forces were only calculated for atoms within 1.1 nm of one another. The temperature was held constant using the velocity rescale thermostat, which is a modification of the Berendsen’s coupling algorithm. Finally, simulation 500 ns long in the NPT ensemble at 300K was performed. All analysis have been performed using the GROMACS tools and post processing using python in-house scripts, available from GitHub page.

Docking calculations between LysB_FL_ and (GlcNAc)5 were performed using HADDOCK version 2.4. We defined the LysB_FL_ active residues 1 to 85, based on Pesto bioinformatic prediction^52^ using protein-ligand approach and on^53^. We defined passive the full 5 GlcNAc ligand. Standard parameters were used during the three different stages (it0, it1 and water refinement), calculating 1,000 structures in it0, of which 200 were refined in it1 and then explicit water.

#### Cell Culture

Cells were maintained at 37°C in a humidified atmosphere containing 5% CO_2_. Media was supplemented with 10% fetal bovine serum (ThermoFisher), 100 U/mL penicillin (ThermoFisher), 0.1 mg/mL streptomycin, and 0.25 μg/mL amphotericin B (ThermoFisher). Max Planck Institute cell-2 (MPI-1)^21,22^ (kindly provided by Marina Freudenberg, Max Planck Institute of Immunology and Epigenetics, Freiburg, Germany) were cultured in Roswell Park Memorial Institute (RPMI) 1640 medium (ThermoFisher) supplemented with 30 ng/mL of murine GM-CSF (Prepotech). MPI-2 cells were passaged twice a week at a concentration of 2 x 10^5^ cells/mL. Cells from passages 8 (p8) to 70 (p70) were used. Cells routinely underwent to Mycoplasma test.

#### MTT viability assay

2 x10^4^ MPI-2 cells/well were seeded in a flat clear-bottom 96 well plates (ThermoFisher) and incubated at 37°C with 5% CO_2_ for 24 hours before the treatment with 15 nmol of either sfGFP or LysB_FL._ After 24 hours of incubation with the recombinant proteins, 10 µl of MTT reagent solution (5 mg/ml) was added to each well and the plate was incubated for 2 hours at 37°C in a CO_2_ incubator. Next, the media with the MTT solution was discarded and 100 µl of solvent solution (HCl 4 mM and 0.1% NP40) was added to each well. The plate was placed on an orbital shaker for 15 minutes at room temperature to dissolve the formazan crystal. At last, the absorbance was measured using a microplate reader at a wavelength of 570 nm.

#### Luminescence viability assay

2 x10^4^ MPI-2 cells/well were seeded in a white clear-bottom 96 well plates (ThermoFisher) and incubated at 37°C with 5% CO_2_ for 24 hours. Then mAMs were either infected or not for 4 hours with *M. tuberculosis* suspension (MOI 1:5). After 4 hours extracellular bacteria were abundantly washed out and the cells were either treated or not with 15 nmol of either sfGFP or LysB_FL._ Over-night. After the overnight incubation with the recombinant proteins, the recombinant proteins were washed out and 100 µl of fresh media supplemented with RealTime-Glo™ MT was added to each well. Viability assay was then performed following the manufacturer’s instruction and luminescence was read as a proxy of cell viability using a plate reader (SPARK, Tecan).

#### Endolysin cellular uptake

4x10^4^ MPI-2 cells/well were seeded in µ-Slide 8 well (IBIDI®; Glass Bottom) and incubated at 37°C with 5% CO_2_ for 24 hours. Then, samples were either treated or not with 15 nmol of sfGFP or LysB_FL_ for 3, 6 and 24 hours. After 24 hours of incubation with the recombinant proteins, MPI-2 cells were fixed by adding 100 μl 4% PFA and incubated at room temperature for 10 minutes. Then the PFA was quickly washed with PBS, and the cells membrane and their nucleus were marked by incubating the µ-Slide with CellMask^TM^ DeepRed (Dilution 1:2000) (Invitrogen^TM^) and Hoechst 35580 (Dilution 1:1000) (Invitrogen^TM^) at room temperature for 10 minutes. The sample was then quickly washed with PBS, and finally stored in PBS at +4°C until sampling.

#### Lipid droplet staining

4x10^4^ MPI-2 cells/well were seeded in µ-Slide 8 well (IBIDI®; Glass Bottom) and incubated at 37°C with 5% CO_2_ for 24 hours. Then, samples were either treated or not with 15 nmol of sfGFP or LysB_FL_ for 3, 6 and 24 hours. After 24 hours of incubation with the recombinant proteins, MPI-2 cells were fixed by adding 100 μl 4% PFA and incubated at room temperature for 10 minutes. Then the PFA was quickly washed with PBS, and the cells were permeabilized with 0.5% Triton^TM^ X-100 (Sigma) and incubated at room temperature for 10 minutes. Then the samples were washed three times in PBS and Nile Red solution (5 μg/mL, final concentration) solution was used to detect the presence of lipid droplet. Cells were incubated with Nile Red solution for 30 minutes and then washed three times with PBS. Finally cellular membrane and their nucleus were marked by incubating the µ-Slide with CellMask^TM^ DeepRed (Dilution 1:2000) (Invitrogen^TM^) and Hoechst 35580 (Dilution 1:1000) (Invitrogen^TM^) at room temperature for 10 minutes. The sample was then quickly washed with PBS, and finally stored in PBS at +4°C until sampling.

#### Gene expression analysis

4 x 10^5^ MPI-2 were seeded in a 32 mm dish and incubated at 37°C with 5% CO_2_ for 24 hours. Then, samples were either treated or not with 15 nmol of sfGFP or LysB_FL_ for 24 hours. At each selected time point, RNA extraction was performed using Monarch® Total RNA Miniprep kit (NEB) following the manufacturer’s instruction. Total RNA was eluted in 30 µl, quantified at ThemoScientificTM NanoDropTM One/OneC Microvolume UV-Vis Spectrophotometer, and stored at −80°C.

cDNA was generated starting from 150 ng of RNA using ProtoScript® II First Strand cDNA Synthesis Kit (NEB) using random hexamers in agreement with the manufacturer’s instruction. The murine primers used are indicated in Key Resource Table. qRT-PCRs were carried out using SensiFAST SYBR Low-ROX Kit (Aurogene), 0.3 µM primers, and 1 µL cDNA diluted 1:4. *Gapdh*-relative quantification was run on Quant Studio Real-Time PCR System (ThermoFisher). The gene expression for each technical triplicate was measured using the 2_-ΔCt_ method.

#### mAMs Infection

##### Gene expression analysis

4 x 10^5^ MPI-2 were seeded in a 32 mm dish and incubated at 37°C with 5% CO_2_ for 24 hours. Then, samples were infected using an MOI 1:5 of a AGT1 suspensions for 4 hours. 4 hours post infection the cells were extensively washed with PBS and treated or not with either 15 nmol of LysB_FL_ or sfGFP overnight. The following day RNA extraction was performed as previously described.

##### Confocal microscopy

4 x 10_4_ MPI-2 cells/well were seeded in µ-Slide 8 well (IBIDI®; Glass Bottom) and incubated at 37°C with 5% CO_2_ for 24 hours. Then, samples were infected using an MOI 1:5 of a AGT1 suspensions for 4 hours. 4 hours post infection the cells were extensively washed with PBS and treated or not with either 15 nmol of LysB_FL_ or sfGFP overnight. The following day, the cells were washed with PBS and then fixed with 4% PFA (ChemoCruz®) for 20 minutes at room temperature. Finally cellular membrane and their nucleus were marked by incubating the µ-Slide with CellMask^TM^ DeepRed (Dilution 1:2000) (Invitrogen^TM^) and Hoechst 35580 (Dilution 1:1000) (Invitrogen^TM^) at room temperature for 10 minutes. The sample was then quickly washed with PBS, and finally stored in PBS at +4°C until sampling.

#### High-resolution confocal microscopy

Fluorescence snapshot imaging was acquired using an inverted STELLARIS 8 Confocal Microscope equipped with a 410 to 850 tunable pulsed wave white laser. All the samples for imaging were prepared by dispensing 0.5 μl of batch culture between two #1.5 coverslips. Exponential-phase *M. smegmatis* RFP reported strain was either treated or not with 15 nmol of LysB_FL_ or sfGFP for 3 hours. Exposure condition: tdTomato and sfGFP were acquired by imaging the sample at 575 nm and 485 nm.

Exponential-phase *M. tuberculosis* RFP reported strain was either treated or not with 15 nmol of LysB_FL_ or sfGFP for 3 hours. Then, the samples were fixed ON with 4% PFA at 4°C. Exposure condition: tdTomato and sfGFP were acquired by imaging the sample at 575 nm and 485 nm.

#### QUANTIFICATION AND STATISTICAL ANALYSIS

##### Image analysis

Image analysis and segmentation were performed using NIS-Elements v. 5.41. Nucleus were identified by modulating DAPI intensity threshold, circularity, and dimension to exclude unfocused objects or artifacts. The cell area was defined by creating a second binary mask using the growing function from the DAPI channel to the CellMask channel. Also, in this case a CellMask threshold intensity was set. Cells touching the field of view border were further excluded from the analysis. AGT1 segmentation was performed using the bright-spot tools. To optimize AGT1 detection, median filter and local contrast parameters were optimized.

#### Mass Spectrometry data processing

Raw data were analyzed using Proteome Discoverer 2.5 using unclassified bacterial viruses (sp_tr_incl_isoforms TaxID=12333_and_subtaxonomy (v2022-01-30) e PD_Contaminants_2015_5.fasta as search database. The MS/MS spectra were searched against the UniProt Ms6 LysB sequence (UniProt entry Q9FZR9). All searches were performed choosing trypsin as specific enzyme with a maximum number of two missed cleavages. Possible modifications included carbamidomethylation (Cys, fixed), oxidation (Met, variable), N-terminal acetylation (variable). The mass tolerance in MS was set to 20 ppm for the first search then 4.5 ppm for the main search and 20 ppm for the MS/MS. Maximum peptide charge was set to seven and seven amino acids were required as minimum peptide length. The “match between runs” feature was applied for samples having the same experimental condition with a maximal retention time window of 0.7 minute. One unique peptide to the protein group was required for the protein identification. A false discovery rate (FDR) cutoff of 1 % was applied at the peptide and protein levels.

#### Statistic

Plots and statistical analyses were generated using Prism 10 (GraphPad Software), FlowJo. Two-way ANOVA, followed by correction for multiple comparisons, was performed to evaluate statistical significance between multiple groups. Brown-Forsythe, Welch, and one-way ANOVA were computed to compare the variation of a single parameter over multiple groups. Plots merge datasets deriving from at least two independent replicates. Significant *p-*values, sample size, and statistical tests are reported in the legends.

## Key Resource Table

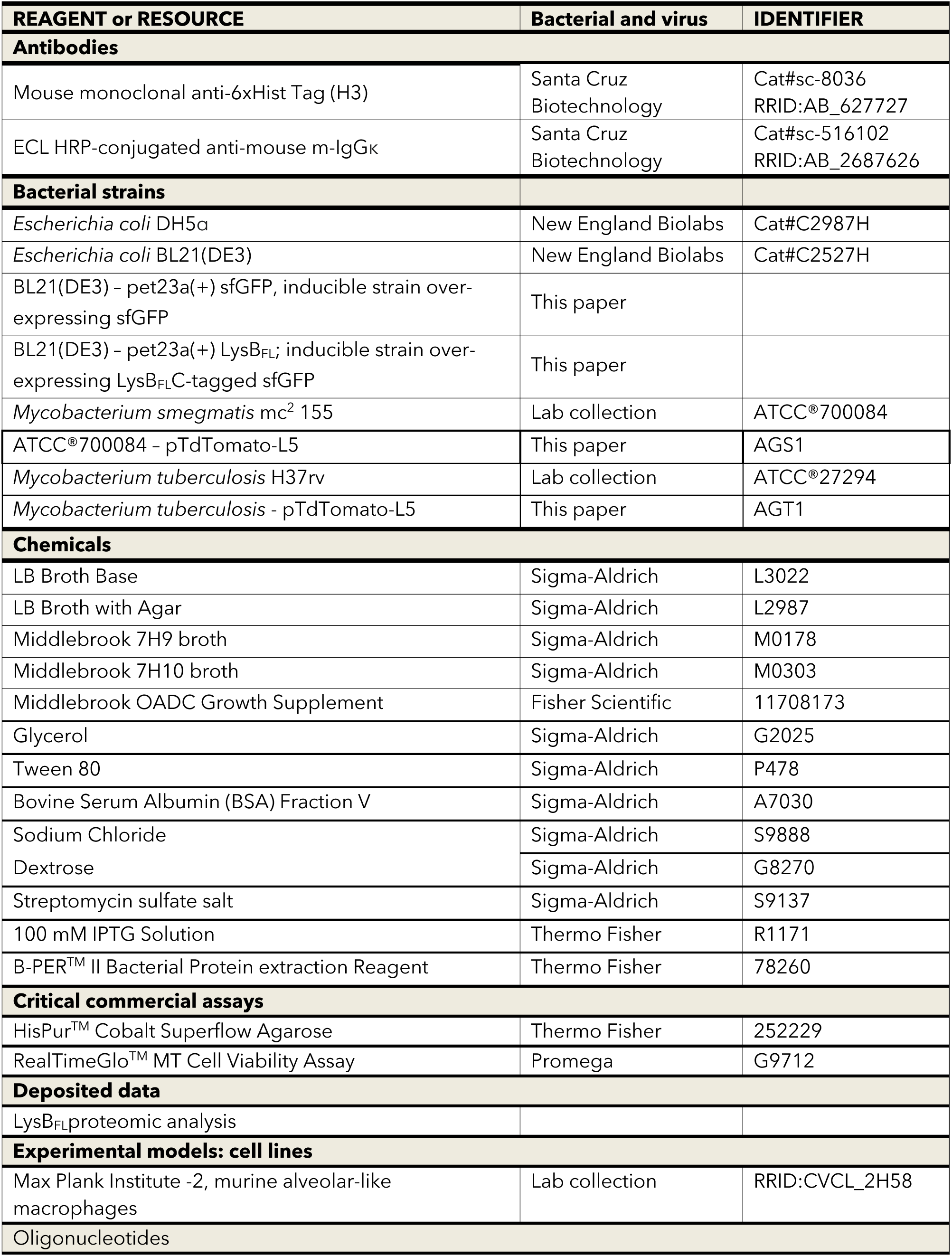
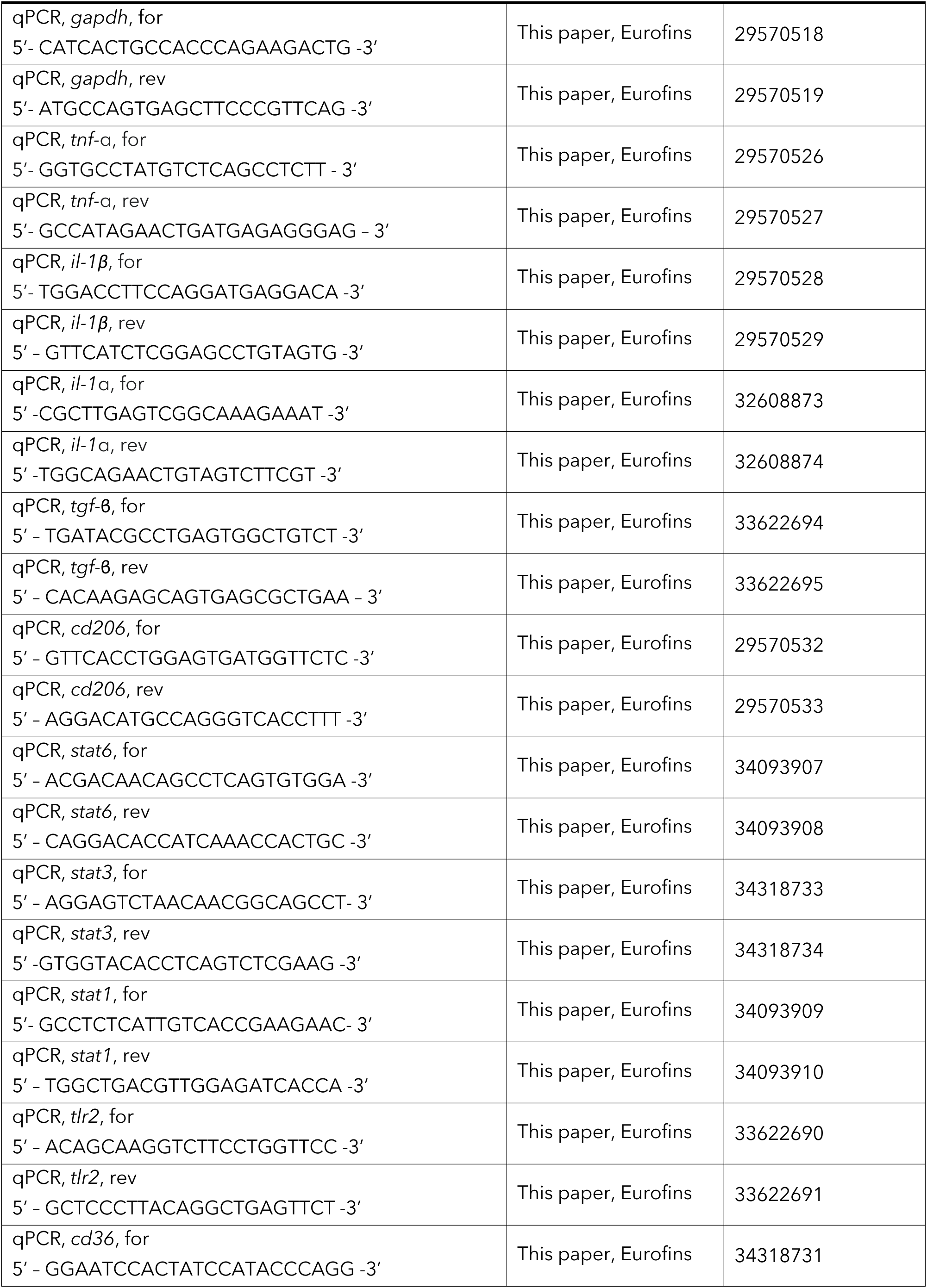
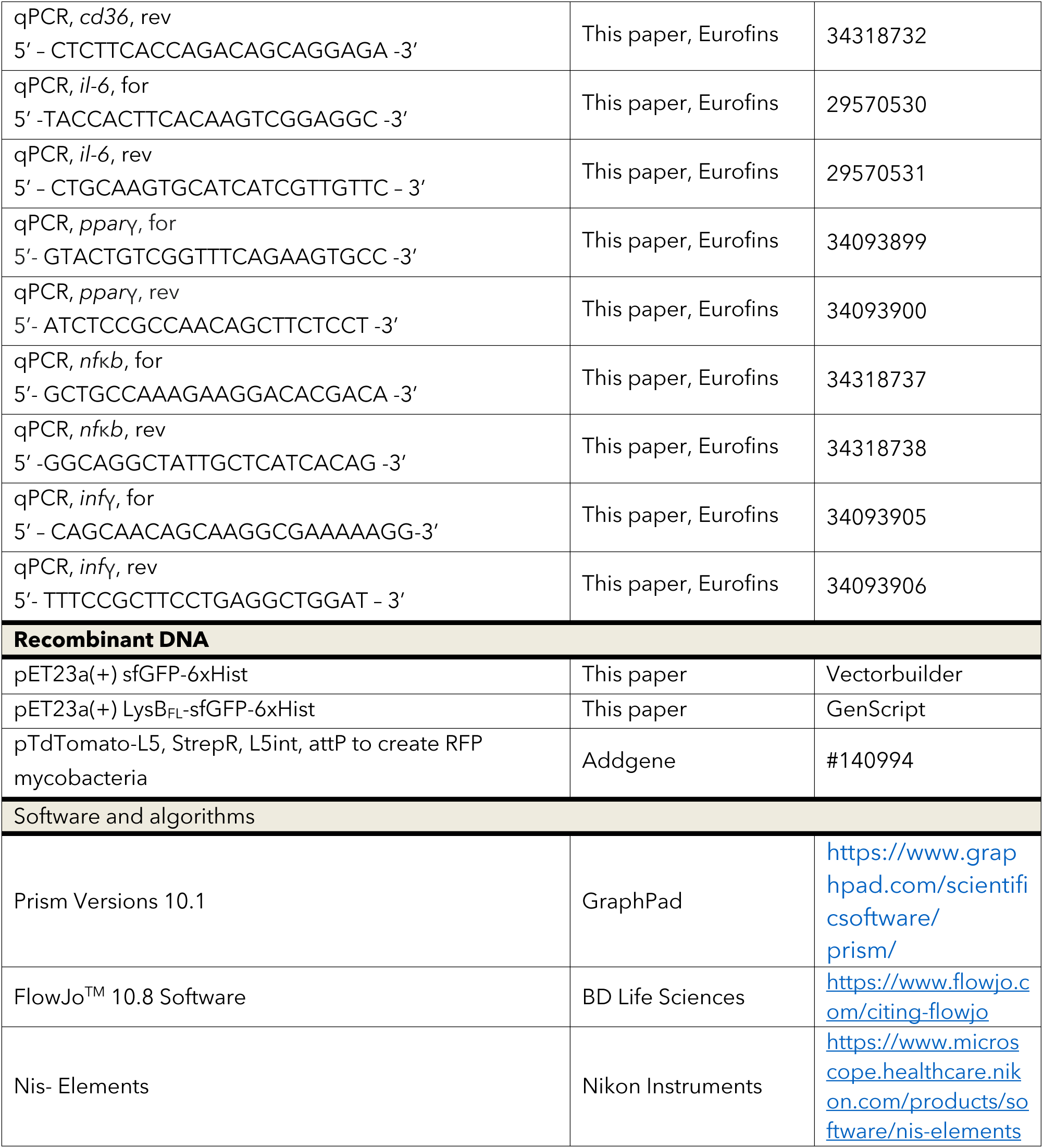

