## Supplementary Material for "Endolysin B as a new archetype in *M. tuberculosis* treatment"

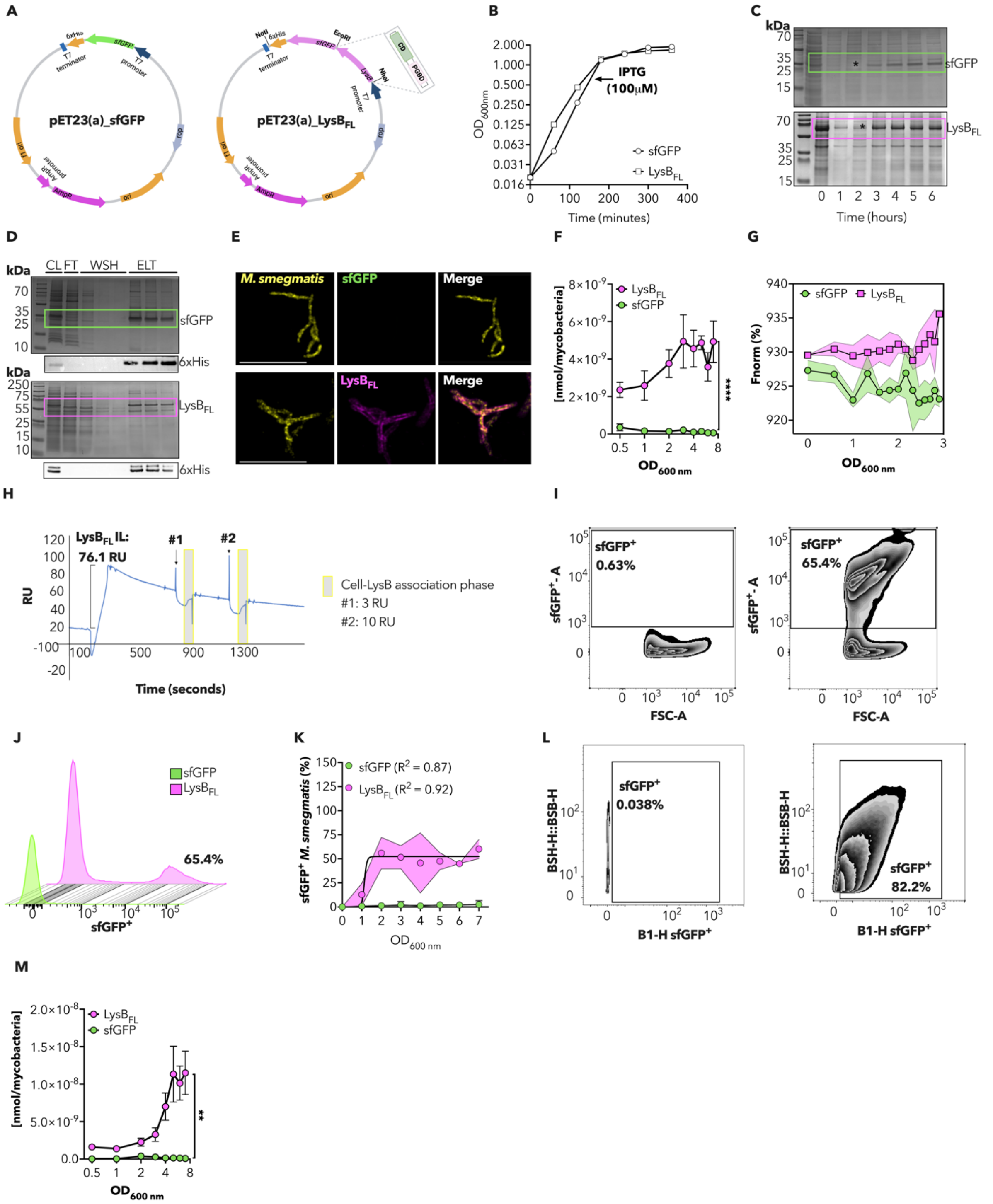

***Figure S1. Recombinant LysB$_{FL}$ bind to M. tuberculosis and M. smegmatis mycomembrane.***

***(A)*** *Schematic representation of the sfGFP (left panel) and LysB$_{FL}$ (right panel) vectors used. Ms6 Endolysin B (LysB$_{FL}$) was C-fused with sfGFP. Tagged LysB$_{FL}$ or sfGFP alone were inserted into the pER23a(+) inducible bacterial expression system. Both constructs were finally inserted in frame with a 6xHist tag.*

***(B)*** *Growth kinetics indicating IPTG-induced over expression of LysB$_{FL}$ (white square) and sfGFP (white circle). The arrow indicates IPTG addition (100 µM final concentration).*

***(C)*** *Coomassie showing LysB$_{FL}$ (64 kDa, magenta square) and sfGFP (27 kDa, green square) over-expression.*
***(D)*** *Coomassie (upper panel) and 6xHist immunoblotting (lower panel) validating LysB$_{FL}$ (64 kDa, magenta square) and sfGFP (27 kDa, green square) purification (CL: cellular lysate, FT: flowthrought, WSH: washing, ELT: elution).*
***(E)*** *Representative images of M. smegmatis treated for 3 hours with 15 nmol of either sfGFP (upper panel) or LysB$_{FL}$ (lower panel). M. smegmatis (yellow), sfGFP (green), LysB$_{FL}$ (magenta), are merged. Scale bar 5 µm.*
***(F)*** *Quantification of the total nmol of LysB$_{FL}$ (magenta) and sfGFP (green) bound to M. smegmatis measured after 3 hours incubation with constant concentration of protein (15 nmol) and decreasing concentration of bacteria plotted on a Log2 scale. Mean values ± SD (n=4) are reported. Asterisks denote significant difference by unpaired t-test with Welch's correction: ** P = 0.0051.*
***(G)*** *MST quantification of sfGFP (green) or LysB$_{FL}$ (magenta) binding to M. smegmatis in function of bacterial density expressed in OD$_{600\ nm}$. All samples were incubated with a constant concentration of the two protein (1.25 nmol) for 3 hours and then measured. Solid black lines represent mean values, green and magenta shading indicates SD as reported in the legend. The data shown are from 4 independent experiment.*
***(H)*** *SPR sensorgram corresponding to manual injection of two bacterial samples flowed on LysB$_{FL}$ protein immobilized onto a CM5 sensor chip by Anti-His antibody coupling. The sensorgram represents the reference-subtracted signal of the cells-LysB$_{FL}$ interaction. The two bacterial samples are indicated with #1 and #2, corresponding to OD $_{600}$ 0.3 and OD $_{600}$ 0.8, respectively. The grey bars highlight the association-related response measured as resonance units (RUs) increasing signal.*
***(I)*** *Density plot summarizing sfGFP$^+$ of M. smegmatis with respect to the physical parameter measured by flow cytometry for sample treated with 15 nmol of sfGFP (left panel) or LysB$_{FL}$ (right panel). Positivity was considered for cells with fluorescence value above $10^3$ values.*
***(J)*** *Histogram showing flow cytometry sfGFP$^+$ profile detected in the sample with OD$_{600}$ = 6 of bacteria treated with 15 nmol of either sfGFP (green) or LysB$_{FL}$ (magenta) for 3 hours*
***(K)*** *Histogram quantifying sfGFP$^+$ M. smegmatis bacteria by flow cytometry in samples treated for 3 hours with 15 nmol of either sfGFP (green) and LysB$_{FL}$ (magenta) Black lines indicate mean value ± SD (n=3). Black lines indicate a Sigmoidal 4PL fitting curve. Magenta shading area denote SD value. The data reported are from three independent experiment.*
***(L)*** *Density plot summarizing sfGFP$^+$ of M. tuberculosis with respect to the physical parameter measured by flow cytometry for sample treated with 15 nmol of sfGFP (left panel) or LysB$_{FL}$ (right panel).*
***(M)*** *Quantification of the total nmol of LysB$_{FL}$ (magenta) and sfGFP (green) bound to M. tuberculosis measured after 3 hours incubation with constant concentration of protein (15 nmol) and decreasing concentration of bacteria plotted on a Log2 scale. Mean values ± SD (n=4) are reported. Asterisks denote significant difference by unpaired t-test with Welch's correction: * P = 0.027.*

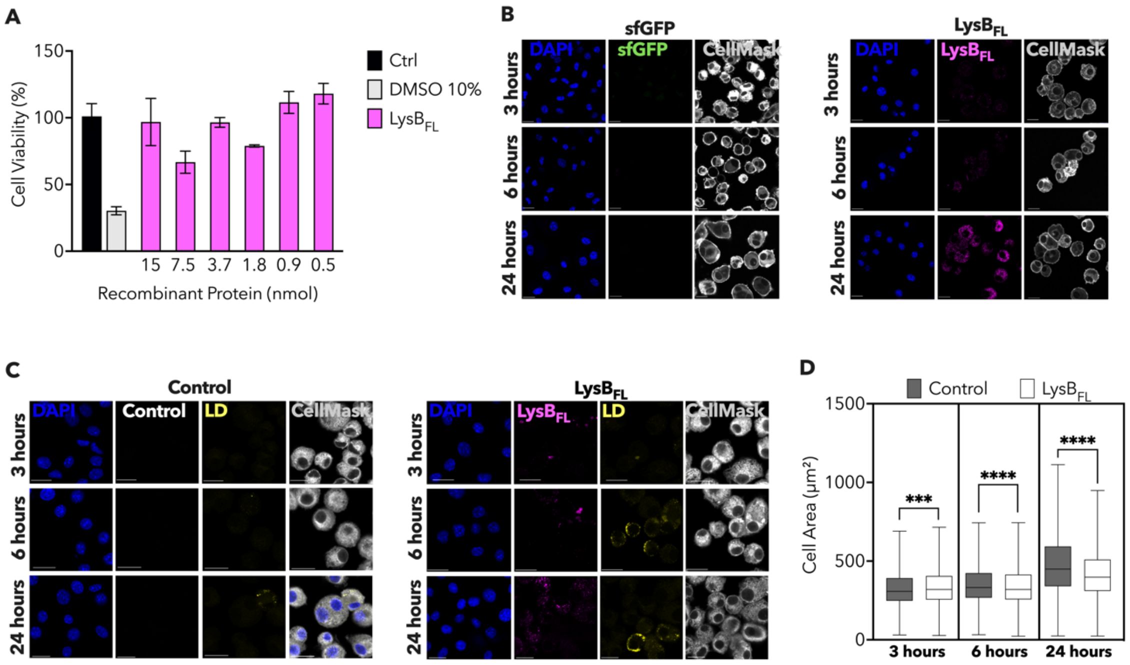


***Figure S2. Cell viability and phenotypic remodulation in AMs-like cells treated with either sfGFP or LysB$_{FL}$ over time.***

***(A)*** *Histogram reporting AMs-like cells viability (MTT) after 24 hours of cells either treated or not with 15 nmol of LysB$_{FL.}$ Black lines indicate mean value ± SD (n=4).*

***(B)*** *Representative images of sfGFP (left panel) and LysB$_{FL}$ (right panel) uptake kinetics in murine AMs-like cells. AMs-like cells (gray), sfGFP (green), LysB$_{FL}$ (magenta), and DAPI (blue) single channels are indicated on the legend. Scale bar 20 µm.*

***(C)*** *Representative images of LD formation in murine AMs-like cells treated (left panel) or not (right panel) with either 15 nmol of r LysB$_{FL}$ for 3, 6 and 24 hours. AMs-like cells (gray), sfGFP (green), LysB$_{FL}$ (magenta), DAPI (blue), LD (yellow) single channels are indicated on the legend. Scale bar 20 µm.*

***(D)*** *Box plot quantifying AMs-like cell area after treatment with 15 nmol of LysB$_{FL}$ (white) or in steady-state (gray) over time. Black lines indicate mean value ± SD (37 > n < 2050). Asterisks denote significant differences by One-way ANOVA followed by Turkey's multiple comparison test: ***p = 0.001, ****p< 0.0001.*

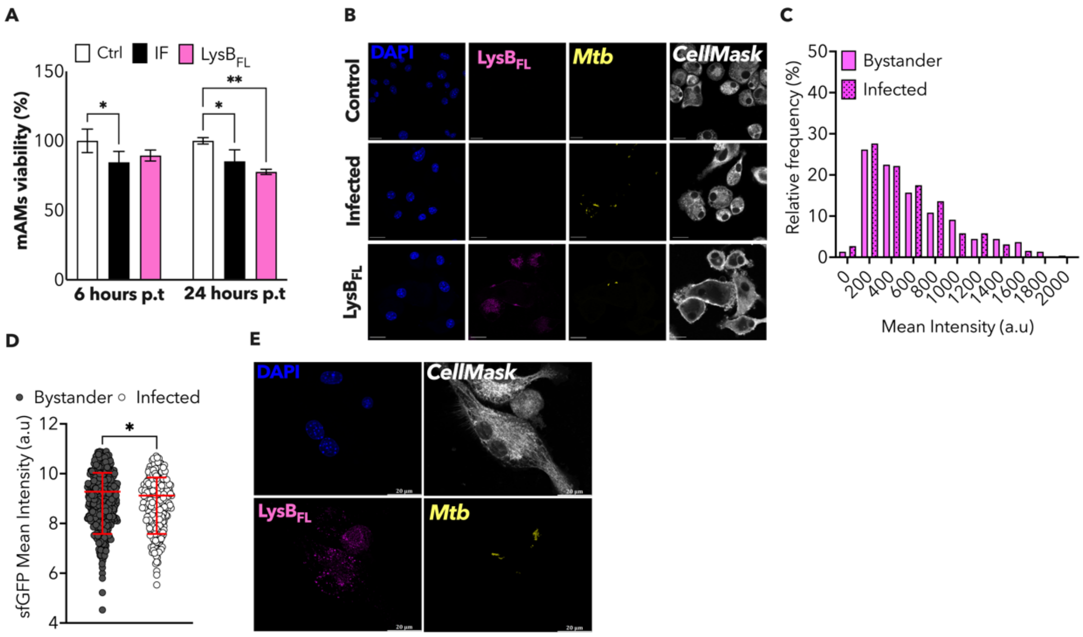


***Figure S3. LysB$_{FL}$ possibly enhance M. tuberculosis control in mAMs.***

***(A)*** *Histogram reporting AMs cells viability detected by luminescence after 6 and 24 hours post mAMs infection with M. tuberculosis for 4 hours, either treated or not with 15 nmol of LysB$_{FL}$. Data are shown from 3 independent experiments. Mean value ± SD are plotted (n=4). Asterisks denote significant differences by Two-way ANOVA followed by uncorrected Fisher's LSD multiple comparison test: *p = 0.02, **p = 0.002.*

***(B)*** *Representative images of mAMs infected (Infected) or not (control) with M. tuberculosis and then treated or not with 15 nmol of LysB$_{FL}$ overnight. AMs cells (gray), LysB$_{FL}$ (magenta), M. tuberculosis (yellow), and DAPI (blue) are indicate in the legend. Scale bar 20 µm.*

***(C)*** *Histogram showing LysB$_{FL}$ distribution in either bystander (plain magenta) or infected (dotted magenta) mAMs infected with M. tuberculosis for four hours and then treated ON with 15 nmol of LysB$_{FL}$. Data are shown from four independent experiments.*

***(D)*** *Single mAMs area in bystander cells (black) and infected cells (white). Red lines indicate mean value ± SD (37 > n < 2050). Black lines indicate mean value ± SD (270 > n < 550). Asterisks denote significant differences by unpaired t-test with Welch's correction: **P = 0.0023.*

***(E)*** *Maximum projection of mAMs infected with M. tuberculosis for four hours and then treated with 15 nmol of LysB$_{FL}$ overnight. DAPI in blue, mAMs in gray, LysB$_{FL}$ in magenta e M. tuberculosis in yellow. Scale bar 20 µm.*
